# Gut Microbiota Dysbiosis Drives Myocardial Hypertrophy Through GBP2b/GBP1-Mediated Immune Reprogramming and Exosomal Signaling in Chronic Colitis

**DOI:** 10.64898/2026.05.27.728214

**Authors:** Yadong Wang, Jian Li, Junqing An, Vu L. Ngo, Shuhan Wang, Zhenkai Hao, Chao Li, Hirohito Abo, Ye Ding, Jun Zou

**Affiliations:** Center for Inflammation, Immunity and Infection, Institute for Biomedical Sciences, Georgia State University, Atlanta, GA, USA

**Keywords:** gut dysbiosis, myocardial hypertrophy, guanylate-binding protein 2b, immunometabolism

## Abstract

**BACKGROUND:** Patients with inflammatory bowel disease (IBD) are at increased risk of cardiovascular disease, yet the mechanisms linking chronic intestinal inflammation to cardiac dysfunction remain poorly understood. IBD is characterized by profound gut microbiota dysbiosis, which we hypothesize drives systemic immune dysregulation and contributes to cardiac dysfunction.

**METHODS:** A chronic colitis mouse model was used to assess gut microbiota dysbiosis, systemic immune cell metabolism, and cardiac remodeling. Cardiac outcomes were evaluated by echocardiography, histology, and molecular analyses. Mechanisms were examined using fecal microbiota transplantation, immune cell depletion, exosome transfer, bone marrow chimeras, RNA-seq, co-immunoprecipitation, confocal microscopy, and siRNA-mediated gene silencing.

**RESULTS:** Chronic DSS colitis induced cardiac dysfunction, hypertrophy, and fibrosis in mice. These changes were accompanied by sustained gut microbiota dysbiosis, metabolic reprogramming, and mitochondrial dysfunction in circulating immune cells. Fecal microbiota transfer experiments demonstrated that colitis-associated microbiota were sufficient to reprogram systemic immune cells and promote cardiac dysfunction. Immune cell depletion studies identified macrophages as key mediators of colitis-associated cardiac injury. Colitis increased systemic lipopolysaccharide (LPS) translocation, bone marrow chimera experiments demonstrated that hematopoietic TLR4 signaling was required for immune cell metabolic remodeling and cardiac dysfunction during chronic colitis. Transcriptomic analysis identified guanylate-binding protein 2b (GBP2b/GBP1, hereafter referred to as GBP1) as a key downstream effector of LPS-TLR4 signaling. Upon LPS stimulation, GBP1 localized to mitochondria, where it interacted with DRP1 and FIS1 to promote mitochondrial fission, oxidative stress, and enhanced immune cell migration into the heart. In addition, GBP1 was secreted via exosomes, which were taken up by cardiomyocytes and contributed to hypertrophic remodeling, and cardiac dysfunction.

**CONCLUSIONS:** These findings establish the LPS-TLR4-GBP1 axis as a key driver of colitis-associated cardiovascular dysfunction and highlight this pathway as a promising therapeutic target for reducing cardiovascular risk in patients with IBD.

**Novelty and Significance:** *What Is Known?:*

- Patients with inflammatory bowel disease have an increased risk of cardiovascular dysfunction that cannot be fully explained by traditional cardiovascular risk factors.
- Gut microbiota dysbiosis and chronic innate immune activation are hallmarks of inflammatory bowel disease, but their direct contribution to cardiac remodeling remains unclear.

*What New Information Does This Article Contribute?:*

- Chronic colitis-associated gut microbiota dysbiosis induces systemic immune cell metabolic and mitochondrial reprogramming that is sufficient to drive cardiomyocyte hypertrophy and cardiac dysfunction.
- Hematopoietic Toll-like receptor 4 signaling links colitis associated gut microbiota to immune metabolic dysfunction and cardiac impairment, establishing a causal gut-immune-heart axis.
- Guanylate-binding protein 2b (GBP2b/GBP1) is identified as a critical downstream effector that promotes mitochondrial fission, oxidative stress, immune cell cardiac infiltration, and exosome-mediated cardiac remodeling.

## INTRODUCTION

Cardiovascular disease (CVD) remains a leading cause of global morbidity and mortality.^1–3^ Epidemiological studies indicate that patients with inflammatory bowel disease (IBD) exhibit a significantly increased risk of heart failure, myocardial infarction, and other forms of cardiac dysfunction.^4–6^ Notably, Large population-based studies have shown that this elevated cardiovascular risk occurs despite a lower prevalence of traditional CVD risk factors in IBD patients, with a∼26-59% higher incidence of coronary heart disease compared with non-IBD controls.^7,8^ IBD is marked by chronic intestinal inflammation, epithelial barrier disruption, and increased permeability, enabling luminal antigens and microbial products to enter the circulation and exert systemic effects.^9,10^ Thus, IBD is increasingly recognized as a systemic inflammatory disorder with important cardiovascular implications.^11^ However, the mechanistic link between chronic intestinal inflammation and cardiac dysfunction remains poorly understood.

Gut microbiota dysbiosis are increasingly recognized as an important contributor to cardiovascular pathology.^12^ During dysbiosis, alterations in microbial structural components and metabolites, such as peptidoglycans, trimethylamine N-oxide (TMAO), short-chain fatty acids, and bile acids, can modulate host immunity and cardiovascular function.^13^ Dysbiosis is a hallmark of IBD and is characterized by loss of beneficial commensals and expansion of proinflammatory or pathogenic taxa, further compromising the epithelial barrier and promoting translocation of microbial products that drive inflammation.^14–16^ Guanylate-binding protein 2b (GBP2b) in mice and its human ortholog GBP1 are inflammation-inducible GTPases, particularly responsive to IFN-γ, and have traditionally been studied for their antimicrobial functions.^17,18^ Growing evidence indicates that GBP1 also regulates inflammatory signaling and contributes to autoimmune diseases and cancer.^19,20^ However, whether and how GBP1 influence cardiovascular pathology remains unknown.

Emerging evidence suggests that immune cell metabolism and mitochondrial dynamics are critical determinants of inflammatory cell function and tissue infiltration.^21–23^ Mitochondrial fission and fusion govern not only bioenergetic flexibility but also reactive oxygen species generation, cytokine production, and migratory capacity of innate immune cells.^24–27^ Dysregulated mitochondrial dynamics in immune cells have been implicated in chronic inflammatory diseases and cardiovascular remodeling;^28,29^ however, the molecular effectors linking inflammatory signals to immune cell mitochondrial remodeling and subsequent cardiac injury have not been defined.

Using a chronic colitis mouse model, we show that colitis-associated dysbiosis, particularly increased exposure to the bacterial component lipopolysaccharide (LPS), drives systemic immune cell metabolic reprogramming and contributes to cardiac dysfunction. We further demonstrate that LPS-induced GBP1 promotes mitochondrial fission and metabolic rewiring in immune cells, enhancing their infiltration into the heart. In addition, GBP1 is enriched in circulating exosomes during colitis, and these exosomes can be taken up by cardiomyocytes, where they may contribute to hypertrophy remodeling and functional impairment. Together, our findings reveal a dual role for GBP1, acting intracellularly to regulate mitochondrial dynamics and immunometabolism, and extracellularly *via* exosomes to mediate microbiota-driven inflammatory signaling that promotes cardiac dysfunction.

## METHODS

### Animal Models and Treatments

C57BL/6 wild-type (WT), Rag1 knockout (Rag1 KO), and Toll-like Receptor 4 knockout (TLR4 KO) mice were obtained from Jackson Laboratory and bred at Georgia State University. At 10 weeks of age, or as specified, mice were fed a grain-based chow diet and subjected to chronic colitis induction with 1% DSS in drinking water (Lot #S7102, MP Biomedicals; MW 36,000-50,000) for one week, followed by one week of regular water, for a total of seven weeks, or as specified. Control mice received regular water only. All procedures were approved by the Georgia State University IACUC (Protocol #A24025). Body weight was monitored regularly. At study termination, epididymal fat mass, spleen, liver, kidney, muscle, colon and heart were collected for subsequent analyses. Indirect calorimetry analysis was performed using a PhenoMaster system (TSE Systems, Bad Homburg, Germany) to assess food intake, water intake, locomotor activity, oxygen consumption, carbon dioxide production, respiratory exchange ratio, and energy expenditure, as previously described. ^30^

### Hematology and Flow Cytometry Analysis

Blood was collected from mice into heparinized tubes, and the immune cell composition was analyzed using a hematology analyzer (VetScan HM5; Abaxis, CA, USA). Colon lamina propria cells and heart-infiltrating immune cells were isolated as previously described.^31^ Non-specific binding was blocked with 10 µg/ml of anti-CD16/ CD32 (Bio X Cell, BP0307) for 10 minutes in FACS buffer. For surface staining, cells were suspended in 0.1 ml of FACS buffer on ice and incubated for 20 minutes with fluorochrome-conjugated antibodies targeting CD45 (Biolegend, 103155), MHC-II (BD Biosciences, BDB563415), CD11b (Biolegend, 101225), Ly6C(Invitrogen, 45-5932-82), CD11C (BD Biosciences, BDB563735), CD301(Biolegend, 145707), F4/80 (Biolegend, 123114), CD64 (Biolegend, 139309), and Ly6G(Biolegend, 127622), to identify innate immune cell populations. After three washes, the stained cells were analyzed using a flow cytometer. Bone marrow cells were collected by flushing with cold PBS, filtered through a 70-μm cell strainer, and treated to remove red blood cells. Hematopoietic stem and progenitor cells were labeled with antibodies against CD127(Biolegend, 12114), Lineage markers (STEMCELL, 19856), c-Kit (Biolegend, 105827), and Sca-1(Biolegend, 108137). Mitochondrial mass, membrane potential, and mitochondrial superoxide were assessed using MitoTracker Green (Invitrogen, M7514), MitoTracker Red (Invitrogen, M36008), and MitoSOX (Invitrogen, M7512), respectively, as previously described.^32^ Flow cytometry data were processed and analyzed with FlowJo software (TreeStar, Ashland, OR).

### Gut Microbiota Composition Analysis

Gut microbial community structure was characterized using 16S rRNA gene amplicon sequencing according to the Illumina 16S Metagenomic Sequencing Library Preparation protocol. Briefly, total genomic DNA was extracted and used for PCR amplification of either the V4 hypervariable region of the 16S rRNA gene using primers 515FB (5 ′ -TCGTCGGCAGCGTCAGATGTGTATAAGAGACAGGTGYCAGCMGCCGCGGTAA-3 ′) and 806RB (5 ′ GTCTCGTGGGCTCGGAGATGTGTATAAGAGACAGGGACTACNVGGGTWTCTAAT-3), or the V3 – V4 region using primers 341F (5 ′ -TCGTCGGCAGCGTCAGATGTGTATAAGAGACAGCCTACGGGNGGCWGCAG-3 ′) and 805R (5 ′ - GTCTCGTGGGCTCGGAGATGTGTATAAGAGACAGGACTACHVGGGTATCTAATCC-3). PCR amplicons were purified with Ampure XP magnetic beads. A second PCR was carried out to incorporate dual indices and Illumina sequencing adapters with the Nextera XT Index Kit. Libraries were quantified, normalized, pooled at equimolar concentrations, and re-quantified prior to sequencing on an Illumina MiSeq platform with paired-end 2 × 250 bp reads. Sequencing data were demultiplexed and processed using the DADA2 workflow within QIIME 2, including quality filtering, denoising, read merging, and chimera removal. Beta diversity was evaluated using unweighted UniFrac distances and visualized by principal coordinate analysis (PCoA) with the emperor plugin in QIIME 2, whereas alpha diversity metrics, including Faith’s phylogenetic diversity and Pielou’s evenness, were calculated in QIIME 2.

### Echocardiographic Assessment of Cardiac Function

Cardiac structure and function were evaluated by transthoracic echocardiography using a Vevo-3100 high-resolution ultrasound imaging system (FUJIFILM Visual Sonics Inc., Toronto, Canada) equipped with a high-frequency transducer. Mice were lightly anesthetized with isoflurane (1.5% in oxygen) to maintain heart rates above 450 beats per minute throughout image acquisition. Two-dimensional parasternal long- and short-axis views were obtained, followed by M-mode recordings at the level of the papillary muscles. Left ventricular end-diastolic and end-systolic dimensions were measured and left ventricular ejection fraction (LVEF), fractional shortening (FS), and left ventricular internal dimension at its widest (diastole, LVIDd) and narrowest (systole, LVIDs) points were calculated using standard formulas. All measurements were averaged from at least five consecutive cardiac cycles and analyzed by an investigator blinded to group allocation using Vevo Lab 2.1.0 software.

### Exosome Isolation, ExosomeTreatment, and Chronic LPS Infusion in Mice

Exosomes were isolated from mouse serum or conditioned culture medium using differential centrifugation combined with polymer-based precipitation or ultracentrifugation. Briefly, samples were subjected to sequential low-speed centrifugation (10 min at 2,000 × g) steps to remove cells and debris, followed by high-speed centrifugation (30 min at 10,000 × g followed by 90 min at 200,000 × g) to pellet extracellular vesicles. The exosome-enriched fraction was resuspended in sterile phosphate-buffered saline. Particle size distribution and concentration were verified by nanoparticle tracking analysis, and exosomal markers were confirmed by immunoblotting. Purified exosomes were quantified based on total protein concentration and diluted in sterile PBS prior to administration. Mice receive exosomes *via* tail vein injection at a concentration of 30 µg/mouse every 3 days. After 28 days, mice heart function was measured, and tissues were harvested for downstream analyses. Osmotic minipumps were used to chronically deliver LPS in mice. Briefly, 6-week Alzet osmotic minipumps were filled with either PBS or LPS from Escherichia coli (Sigma, St. Louis, MO) and implanted subcutaneously to deliver LPS at a dose of 300 μg/kg/day, as previously described.^33^ After 42 days, cardiac function was assessed, and heart weight was measured.

### BMDM Differentiation, siRNA Transfection, and In Vivo Migration Assay

Bone marrow-derived macrophages (BMDMs) were generated from femoral and tibial bone marrow of CD45.1 wild-type mice. Briefly, bone marrow cells were harvested by flushing with cold PBS, filtered through a 70-μm cell strainer, and subjected to red blood cell lysis. Cells were cultured in DMEM supplemented with 10% fetal bovine serum, 1% penicillin/streptomycin, and macrophage colony-stimulating factor (M-CSF, 25 ng/mL). On day 5, the medium was replaced with fresh medium containing an increased concentration of M-CSF (50 ng/mL). On day 6, ranulocyte-macrophage colony-stimulating factor (GM-CSF, 50 ng/mL) was added, and fully differentiated macrophages were collected on day 7, and electroporated using the SG Cell Line 4D-Nucleofector X Kit (Lonza, V4XC-3024) according to the manufacturer’s instructions, with 50 nM siRNAs (Santa Cruz Biotechnology, sc-41707) or corresponding controls. Following transfection, 1 × 10^6^ cells were adoptively transferred into CD45.2 wild-type recipient mice. Mice were subsequently injected with LPS, and hearts were harvested 12 hours later. Donor-derived (CD45.1) and host-derived (CD45.2) macrophage populations in the hearts were analyzed by flow cytometry (FACS).

### Generation of Bone Marrow Chimeras

Healthy wild-type recipient mice were lethally irradiated with 8.5 Gy using an RS-2000 irradiator (Rad Source Technologies) and subsequently reconstituted via intravenous injection of 2 × 10^6^ bone marrow cells derived from either wild-type or TLR4 knockout (KO) donors to generate BM-WT or BM-TLR4KO chimeric mice, respectively. In separate experiments, recipient mice were reconstitute with bone marrow cells from wild-type control or colitis mice. Following transplantation, mice were housed in sterile cages with access to sterile water for the first two weeks. Experimental procedures were conducted for three- or six-weeks post-reconstitution.

### Fecal Microbiota and Bacterial Transplantation

For fecal microbiota transplantation (FMT), fecal samples were collected from healthy or post-DSS colitis mice at 91 days after colitis induction and resuspended in PBS at a concentration of 100 mg/mL. Eight-week-old germ-free (GF) C57BL/6 male mice (Taconic Biosciences) were colonized via oral gavage with 100 μL of the fecal suspension. Colonized GF mice were maintained in isocages for eight weeks prior to assessment of cardiac function. For *E. coli* colonization, bacterial strains were isolated from patients and cultured overnight in BHI medium. Chimeric BM-WT or BM-T4KO mice generated by bone marrow transplantation were orally administered 2.5 × 10⁸ CFU of *E. coli* per mouse, and cardiac function was evaluated after four weeks of colonization.

### Neutrophils and macrophage depletion

Wild-type male mice were subjected to chronic colitis induction with 1% DSS in drinking water. For neutrophil depletion, mice received intraperitoneal injections of anti-Ly6G antibody (Bio X Cell, BE0075; initial dose, 200 μg/mouse; maintenance dose, 100 μg/mouse every other day) for 4 weeks. For macrophage depletion, mice received intraperitoneal injections of anti-CSF1R antibody (Bio X Cell, BE0213; 200 μg/mouse) once weekly, together with intravenous injections of clodronate liposomes (LIPOSOMA; 200 μL/mouse) twice weekly, until day 28 after initiation of DSS treatment.

### Cell Culture, Cell treatment, and Primary Cell Isolation

Circulating immune cells were isolated from whole blood collected into heparin microtubes following brief red blood cell lysis. Briefly, 200 μL of anticoagulated blood was centrifuged, and the plasma was carefully removed. One milliliter of red blood cell (RBC) lysis buffer was then added to the remaining cell pellet and incubated for 5 minutes at room temperature. The sample was subsequently centrifuged, and the cell pellet was washed twice with PBS. The isolated immune cells were finally resuspended in culture medium. Hematopoietic progenitor cells were isolated using the EasySep™ Mouse Hematopoietic Progenitor Cell Isolation Kit (Catalog #19856, STEMCELL Technologies). Cell number and viability were assessed using trypan blue staining and a cell counter

Blood granulocytes and mononuclear cells were separated from heparinized whole blood by using Ficoll-Paque Plus density gradient centrifugation. H9C2 cardiomyocytes were donated by Binghe Wang from Georgia State University and cultured in Dulbecco’s modified Eagle’s medium (DMEM, Corning, 10-013-CV, NY) containing 10% (*v*/*v*) fetal bovine serum (FBS; Sigma-Aldrich, F0926, St Louis, MO) and 1% (*v*/*v*) antibiotics (penicillin and streptomycin; Gibco, 15140-122; Grand Island, NY) in a humidified incubator at 37 °C and 5% CO2. To establish an experimental hypertrophy condition, H9C2 cells were serum starved for 18 h in DMEM containing 1% FBS. In the experiments, H9C2 were co-cultured with blood immune cells isolated from heparinized whole blood following red blood cell lysis, using trans-well chambers with 0.4 µm pore size inserts, or treated with exosomes for 24 h. After treatment, hypertrophic growth of H9C2 cells was assayed by measurement of cell surface area alongside with determination of protein content.

THP1 monocytes were obtained from American Type Culture Collection (ATCC, TIB-202; Manassas, VA) and maintained in RPMI 1640 Medium (Corning, 10-040-CV, NY) supplemented with 10% FBS and 1% antibiotics in a humidified incubator at 37 °C and 5% CO2. For macrophages differentiation, THP1 monocytes (passage 5-10) were seeded at a concentration of 5 × 10^5^ cells/ml in the presence of 200 ng/ml Phorbol 12-myristate 13-acetate (PMA, Sigma Aldrich, P8139, MO) for 24 h, followed by a 48-h rest period in culture media. Raw264.7 macrophages were purchased from ATCC and maintained in RPMI 1640 Medium with 10% FBS and 1% antibiotics in a humidified incubator at 37 °C and 5% CO2.

### Small interfering RNA (siRNA) Transfection

For transfections of GBP1 (Santa Cruz Biotechnology, sc-72088 and sc-41707) in macrophages, differentiated THP1 macrophages or Raw264.7 macrophages were primed by 10 ng/ml LPS for 8 h and were then electroporated using SG Cell Line 4D-Nucleofector X Kit (Lonza, V4XC-3024) according to the manufacturer’s instructions. 50 nM siRNAs as well as their correspondent controls were used for transfections.

### Measurement of Mitochondrial Respiration and Glycolytic Activity

Cellular bioenergetics were assessed using an extracellular flux analyzer (Agilent Technologies) to determine oxygen consumption rate (OCR) and extracellular acidification rate (ECAR). Blood immune cells were seeded into Seahorse XF cell culture microplates at optimized densities following a suspension cells seeding protocol provided by Agilent Technologies. Prior to the assay, cells were washed and incubated in Seahorse assay medium supplemented with glucose (10 mM), pyruvate (1 mM), and glutamine (2 mM), and equilibrated in a non-CO₂ incubator for 1h. For mitochondrial stress tests, OCR was measured at baseline and following sequential injections of 1.5 µM oligomycin, 4 µM FCCP, and 1 µM rotenone/antimycin A. Glycolytic function was assessed by ECAR measurements after sequential addition of 10 mM glucose, 1 µM oligomycin, and 50 mM 2-deoxy-D-glucose. OCR and ECAR values were normalized to cell number.

### Assessment of Glucose, Triglycerides, Cholesterol, LPS, TMAO and ALT

Blood glucose was determined using a Nova Max Plus Glucose Meter and reported in mg/dL. Serum triglyceride and cholesterol concentrations were measured with Infinity Triglyceride Reagent (Thermo Scientific, TR22421) and Cholesterol Reagent (Thermo Scientific, TR13421), respectively, following the manufacturers’ protocols. Levels of lipopolysaccharide (LPS) and trimethylamine N-oxide (TMAO) were quantified using a commercial LPS ELISA kit (MBS2703401) and a Mouse Trimethylamine N-Oxide (TMAO) ELISA kit (BHE13707179), in accordance with the suppliers’ instructions. Serum alanine aminotransferase (ALT) activity was measured using an ALT activity assay kit (E-BC-K235-S, Elabscience).

### MitoTracker Staining and Subcellular Fractionation in THP1 cells

Mitochondrial morphology in THP-1 macrophages was assessed using MitoTracker Green dye. Specifically, Following the indicated treatment, cells were incubated with MitoTracker Green (250 nM) in pre-warmed serum-free medium for 30 minutes at 37 °C in the dark. Fluorescent images were acquired using a laser-scanning confocal microscope (Carl Zeiss Meditec, Inc., Jena, Germany). Mitochondrial fragmentation was quantified by measuring mitochondrial length and calculating the percentage of mitochondria with increased circularity. At least three independent experiments were conducted, and a minimum of 20 cells per condition were analyzed for static morphological assessment.

Subcellular fractionation of THP1 macrophages was carried out with minor adaptations from established methods. Cells were collected and mechanically disrupted by repeated passage through a 27-gauge needle in ice-cold fractionation buffer containing 150 mM KCl, 25 mM Tris-HCl (pH 7.4), 5 mM EDTA, and protease inhibitors. A part of the homogenate was saved as whole cell lysate, and the rest was subjected to low-speed centrifugation (10 min at 800 × g) to pellet nuclei. The post-nuclear supernatant, comprising cytosolic and membrane-associated components, was further processed using a commercial mitochondrial isolation kit (ThermoFisher Scientific, 89874, MA) to enrich for cytoplasm and mitochondrial fractions. Protein concentrations were determined by BCA assay, and equivalent amounts of protein from each fraction were used for subsequent immunoblot analyses.

### Immunoprecipitation

Protein-protein interactions were examined by immunoprecipitation (IP) following standard procedures. Cell lysates were prepared in IP lysis buffer containing Tris-HCl, NaCl, EDTA, and Triton X-100 (pH 7.4). After clarification, total protein concentrations were quantified by BCA assay. Equal amounts of lysate (500 µg) were incubated overnight at 4°C with antibodies targeting DRP1 (Cell Signaling Technology, 8570). Immune complexes were captured, washed repeatedly with lysis buffer to reduce nonspecific binding, and eluted by heating in Laemmli sample buffer. Eluted proteins were subsequently resolved by SDS-PAGE and analyzed by western blotting.

### Immunoblotting Analysis

Total protein extracts from tissues or cultured cells were obtained using RIPA buffer. Protein levels were measured by the BCA method prior to electrophoretic separation by SDS-PAGE. Proteins were transferred onto PVDF membranes, which were then incubated with primary antibodies against ANP (GeneTex, GTX109255), BNP (abcam, A11515), β-MHC (ProteinTech, 22280-1-AP), GBP1 (ProteinTech, 15303-1-AP), DRP1(Cell Signaling Technology, 8570), CD81 (Santa Cruz Biotechnology, sc-9158), GAPDH (Santa Cruz Biotechnology, 32233), or β-actin (Santa Cruz Biotechnology, 47778). Following incubation with HRP-conjugated secondary antibodies, immunoreactive signals were detected using enhanced chemiluminescence. Band intensities were quantified using ImageJ software, and data represent the average values obtained from at least three independent experiments.

### Histology, Immunohistochemistry, and Immunofluorescence staining

Mice heart tissue was excised, fixed overnight in 4% paraformaldehyde, and paraffin embedded. Tissue blocks were sectioned at a thickness of 4 µm and subjected to standard histological staining protocols, including hematoxylin and eosin (H&E) staining for cardiac morphology and Picroirius Red (PSR) staining for collagen visualization. Cardiac wall thickness, cardiomyocyte size, and collagen volume were measured using Image J software.

Paraffin-embedded heart tissue sections were incubated overnight at 4°C with primary antibodies against GBP1 (abcam, ab119236). Antigen-antibody complexes were detected using ImmPRESS HRP Goat anti-Mouse IgG Polymer Kit (Vector, MP-7452-50), followed by color development with DAB substrate (Dako, K3467). Images were acquired using an inverted microscope. Quantification of staining intensity and positive area was performed using ImageJ software.

For immunofluorescence analysis, paraffin-embedded heart tissue sections (after antigen retrieval) or methanol-fixed THP1 macrophages were incubated overnight at 4°C with antibodies against CD68, GBP1, DRP1, FIS1, or appropriate isotype controls. Tissue sections or cells were then incubated with fluorophore-conjugated secondary antibodies at room temperature, and nuclei were counterstained with DAPI. Fluorescent images were captured using a confocal laser scanning microscope. At least three independent experiments were conducted, with a minimum of three high-magnification fields analyzed per experiment. Co-localization between GBP1 and DRP1 (or FIS1) was assessed using Pearson’s correlation coefficient.

### Quantitative Real-Time PCR and RNA Sequencing

Total RNA was isolated from mouse heart tissues, peripheral blood immune cells, or cultured macrophages using TRIzol reagent (Invitrogen, AM9738). For blood immune cells, erythrocytes were removed prior to RNA extraction using red blood cell lysis buffer (ThermoFisher Scientific, 00-4333-57, MA). RNA concentration and purity were determined spectrophotometrically, and equal amounts of RNA were reverse transcribed into complementary DNA using a reverse transcription kit. Quantitative real-time PCR was performed using SYBR Green-based chemistry on a real-time PCR detection system. Relative mRNA levels were calculated using the comparative Ct (2^−ΔΔCt) method, with GAPDH or β-actin serving as the internal reference gene. Primer sequences are provided in the Supplementary Table 1. All reactions were performed in technical triplicates, and experiments were repeated independently at least three times. For RNA sequencing, purified RNA samples were submitted to Novogene for library preparation and high-throughput sequencing. Sequencing read quality was assessed using FASTQC, and downstream RNA-seq data analyses were conducted using the R software platform.

### Serum Fractionation and Functional Screening Assay

To identify the biochemical nature of circulating factors responsible for macrophage activation, serum was collected from control and DSS-induced chronic colitis mice. Samples were subjected to selective biochemical treatments prior to functional testing in macrophages. For heat inactivation, serum aliquots (200 µL) were incubated at 95°C for 10 min, rapidly cooled on ice, and centrifuged at 10,000 × g for 10 min to remove precipitated proteins. Supernatants were collected for downstream assays. For protein digestion, serum was incubated with Proteinase K (final concentration 100 µg/mL) at 55°C for 60 min, followed by enzyme inactivation at 95°C for 10 min. For nucleic acid degradation, serum was treated with DNase I (100 U/mL) and RNase A (50 µg/mL) at 37°C for 60 min. For lipid removal, serum lipids were extracted using a chloroform/methanol phase separation method. The aqueous phase was collected and used for functional assays. For LPS neutralization, serum samples were incubated with polymyxin B (20 µg/mL) at 37°C for 30 min prior to cell treatment. Murine RAW264.7 macrophages were cultured under standard conditions and treated with processed or untreated serum (typically 10% *v*/*v*) for 16 hours. Following stimulation, TNF-α protein levels in culture supernatants were measured by ELISA.

### DiR Labeling and Biodistribution Analysis of Exosomes In *Vivo* and In *Vitro*

Exosomes were isolated from mouse serum by sequential centrifugation followed by ultracentrifugation. Purified exosomes were labeled with the near-infrared lipophilic dye DiR according to the manufacturer’s instructions. Briefly, exosomes were incubated with 5 µM DiR at room temperature for 20 min, followed by ultracentrifugation to remove unbound dye. The exosome pellet was extensively washed with PBS and re-purified by ultracentrifugation to minimize contamination from free dye. For *in vivo* biodistribution analysis, DiR-labeled exosomes were resuspended in sterile PBS and administered to mice via tail vein injection. Control mice received an equivalent amount of free DiR dye processed under identical conditions. At indicated time points, whole-body fluorescence imaging was performed using an in vivo imaging system (IVIS). Mice were subsequently euthanized, and major organs, including liver, lung, spleen, heart, and kidney, were harvested for ex vivo fluorescence imaging. For *in vitro* uptake analysis, H9C2 cardiomyocytes were incubated with DiR-labeled exosomes for 24 h. Live-cell fluorescence images were captured using fluorescence microscopy to evaluate cellular uptake of exosomes.

### Statistical Analysis

All data are presented as mean ± standard error of the mean (S.E.M.). Statistical analyses were performed using GraphPad Prism software. Comparisons between two groups were conducted using unpaired two-tailed Student’s t test. For experiments involving more than two groups, one-way or two-way analysis of variance (ANOVA) was applied, followed by appropriate post hoc tests to correct for multiple comparisons. Normality of data distribution was assessed prior to parametric testing. A *p* value < 0.05 was considered statistically significant. The number of biological replicates and specific statistical tests used are indicated in the corresponding figure legends.

### Data Availability

Data supporting the findings of this study are available from the corresponding author upon reasonable request. Sequencing datasets have been deposited in the NCBI Sequence Read Archive (SRA) under BioProject accession number PRJNA1464720.

## RESULTS

### Chronic DSS-Induced Colitis Drives Cardiac Dysfunction and Hypertrophy in Mice

Patients with IBD exhibit a significantly increased risk of developing heart dysfunction.^34^ To model this clinical association, we induced chronic colitis in mice using low-dose dextran sulfate sodium (DSS, 1%), which disrupts the colonic epithelium and promotes sustained intestinal inflammation. Mice underwent repeated DSS cycles (one week on DSS followed by one week off, as outlined in Figure S1A) and developed hallmark features of chronic colitis, including mild weight loss (Figure S1A), shortened colon length, and an increased colon weight-to-length ratio (Figure S1B and S1C), all indicative of persistent low-grade inflammation. This colonic inflammatory state was further confirmed by flow cytometry analysis (Figure S1D), which showed increased recruitment of neutrophils, inflammatory monocytes and macrophages (Figure S1E-1G), elevated fecal lipocalin-2 (Lcn2) levels (Figure S1H), and characteristic inflammatory changes observed by H&E staining (Figure S1I). To determine whether chronic colitis induces systemic tissue injury, we examined major peripheral organs. Histological analysis revealed no overt pathological alterations in the kidney, liver, lung, skeletal muscle, or epididymal white adipose tissue (eWAT), although mild inflammatory features were observed in the spleen and liver, key sites of systemic immune activity (Figure S2A). Consistent with preserved hepatic integrity, no liver fibrosis was detected, and serum alanine aminotransferase (ALT) activity remained unchanged (Figure S2B), further indicating the absence of overt liver dysfunction.

Strikingly, chronic colitis resulted in significant cardiac dysfunction. Echocardiography revealed reduced left ventricular ejection fraction (LVEF) and fractional shortening (FS), accompanied by increased left ventricular internal diameter at diastole (LVIDd) and systole (LVIDs), in both male and female mice (Figure 1A and 1B; Figure S2C), demonstrating impaired systolic performance. Structural remodeling was further evidenced by multiple indicators of cardiac hypertrophy, including increased heart-to-body weight ratio and tibia length-normalized heart weight (Figure 1C-1E; Figure S2D), greater ventricular wall thickness, and enlarged cardiomyocyte size (Figure 1F-1H). Increased interstitial collagen deposition, indicative of cardiac fibrosis, was also observed (Figure 1F and 1I), marking progression toward maladaptive remodeling. To define the molecular changes underlying these phenotypes, we performed RNA sequencing (RNA-seq) on heart tissue from DSS-treated and control mice. We observed widespread transcriptional alterations (Figure S2E), including significant upregulation of hypertrophy-associated genes (Figure 1J) such as Nppa, Nppb, and Myh7, directly confirming the hypertrophic state at the transcriptomic level. The increased expression of Nppa, Nppb, and Myh7 were validated by quantitative RT-PCR (Figure 1K) and further supported at the protein level by western blot analysis of atrial natriuretic peptide (ANP), brain natriuretic peptide (BNP), and β-myosin heavy chain (β-MHC) (Figure 1L and 1M). Moreover, KEGG pathway analysis of differentially expressed genes identified several enriched pathways (Figure S2F), with mitochondrial ribosome and ribosome biogenesis pathways among the most prominent (Figure S2G), suggesting mitochondrial stress and impaired cellular homeostasis in the hearts of DSS-treated mice. Collectively, these results demonstrate that chronic intestinal inflammation induces systemic inflammatory responses in the absence of overt organ injury, yet is sufficient to drive cardiac dysfunction, hypertrophy, and fibrotic remodeling.

**Figure 1.**
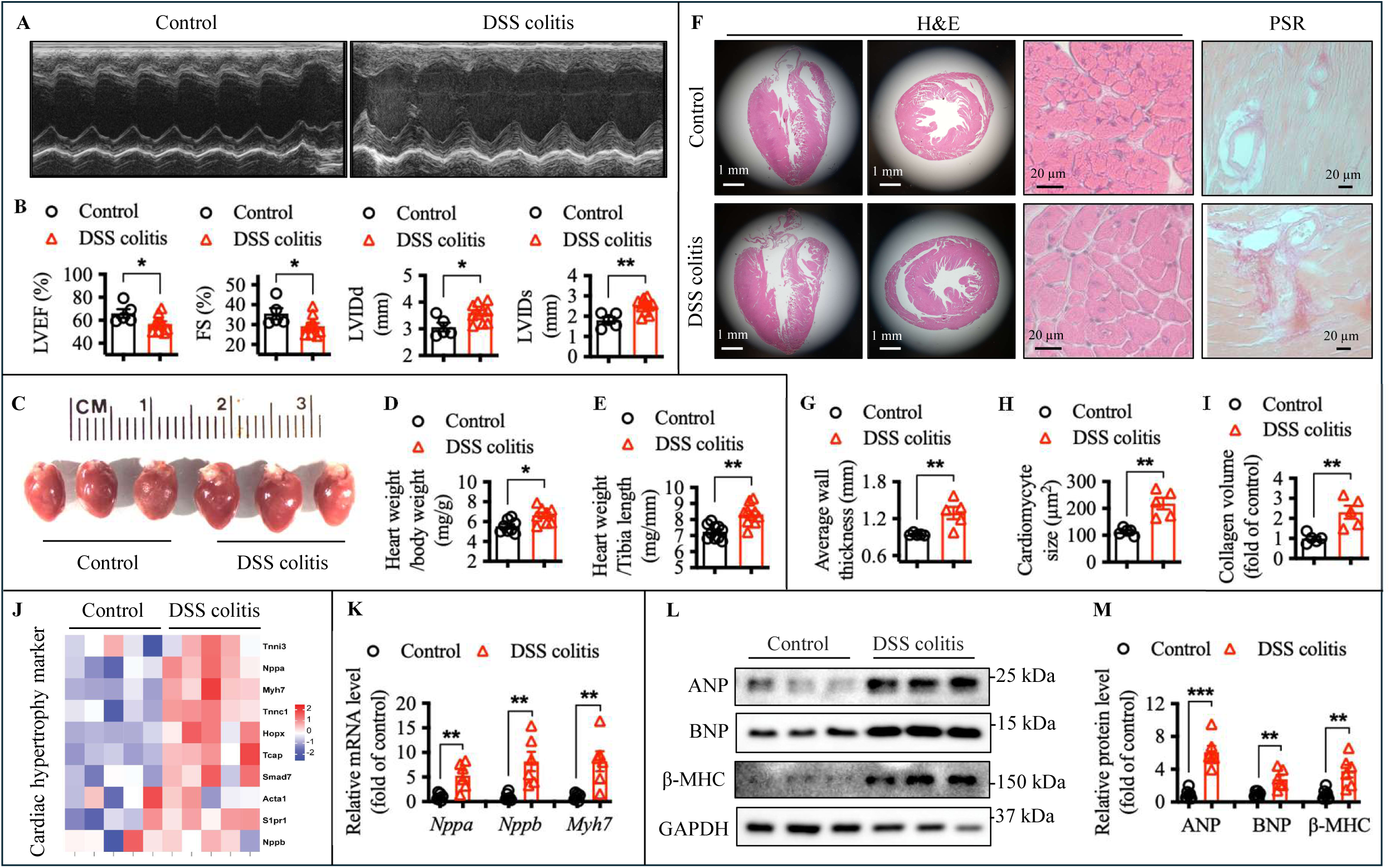
DSS-induced chronic colitis promotes cardiac hypertrophy in mice. **A**, Representative images of echocardiographic analysis in mice treated with vehicle or 1% DSS at day 49 after DSS initiation. **B**, Quantification of left ventricular (LV) ejection fraction (LVEF), fraction shortening (FS), LV end-diastolic internal dimension (LVIDd), and LV end-systolic internal dimension (LVIDs). n=5-10 mice/group. **C**, Representative images of mice heart. **D** and **E**, Quantification of ratios of heart weight to body weight (**D**) and heart weight to tibia length (**E**). n=8 mice/group. **F**, Representative images of hematoxylin-eosin (H&E) and Picrosirius Red (PSR) staining of longitudinal and cross-sections in the heart. **G** through **I**, Quantification of average LV wall thickness (**G**), cardiomyocytes size (**H**), and collagen volume (**I**). n=5 mice/group. **J**, Heatmap of relative mRNA level of the indicated genes in the heart tissue of control or DSS colitis mice. **K** through **M**, Q-PCR (**K**) and Western-blot (**L** and **M**) analysis of cardiac hypertrophy markers, including ANP (gene name: *Nppa*), BNP (gene name: *Nppb*), and β-MHC (gene name: *Myh7*). n=6 mice/group. Data are presented as mean ± S.E.M. with statistical significance indicated as follows: * *p*<0.05, ** *p*<0.01, *** *p*<0.001.

### Colitis-Associated Immune Cells Exhibit Mitochondrial Dysfunction and Proinflammatory Metabolic Reprogramming

To define mechanisms underlying chronic colitis-induced cardiac dysfunction, we first assessed cardiometabolic parameters. Colitis mice exhibited lower blood glucose, while adipose mass, triglyceride, cholesterol, and TMAO levels were unchanged (Figure S3A). Consistently, metabolic cage analysis revealed no differences in caloric intake, respiratory exchange ratio (RER), energy expenditure, or locomotor activity (Figure S3B-S3E), indicating minimal disruption of whole-body metabolic homeostasis, arguing against a primary metabolic contribution to cardiac dysfunction.

Given that colitis-associated systemic immune activation may contribute to CVD, we examined immune cell populations in the circulation and heart during DSS-induced colitis. By day 7, hematology analysis using a hematology cell counter showed increased circulating neutrophils and a modest rise in monocytes, while lymphocyte counts remained unchanged (Figure S4A). To further assess the role of lymphocytes, DSS-induced colitis was performed in Rag1^-/-^ mice (Figure S4B&C), which lack mature T and B cells. Cardiac dysfunction and hypertrophy persisted, as evidenced by reduced LVEF, FS, and increased heart-to-body mass ratio (Figure S4D&S4E), suggesting that lymphocytes are not required and implicating a potential role for innate immune cells. Consistent with this, flow cytometry revealed increased proportions of circulating neutrophils, inflammatory monocytes and macrophage subsets in the blood (Figure 2A-2C). To further characterize these immune cell alterations, we performed RNA-seq analysis of blood-derived immune cells. KEGG analysis of differentially expressed genes revealed that oxidative phosphorylation and reactive oxygen species production were among the top five affected pathways (Figure 2D), indicating profound colitis-associated reprogramming of immune cell metabolism. Consistent with this transcriptomic signature, Seahorse extracellular flux analysis demonstrated that circulating immune cells from day 7 colitis mice exhibited reduced oxygen consumption rate (OCR), and increased extracellular acidification rate (ECAR), reflecting impaired mitochondrial respiration and a compensatory shift toward glycolytic metabolism (Figure 2E&2F). By 42 days after colitis induction, the percentage of circulating neutrophils and inflammatory macrophage subsets remained elevated (Figure S5A), and metabolic dysfunction persisted, as evidenced by sustained reductions in OCR and increased ECAR in blood-derived immune cells (Figure S5C&S5D).

**Figure 2.**
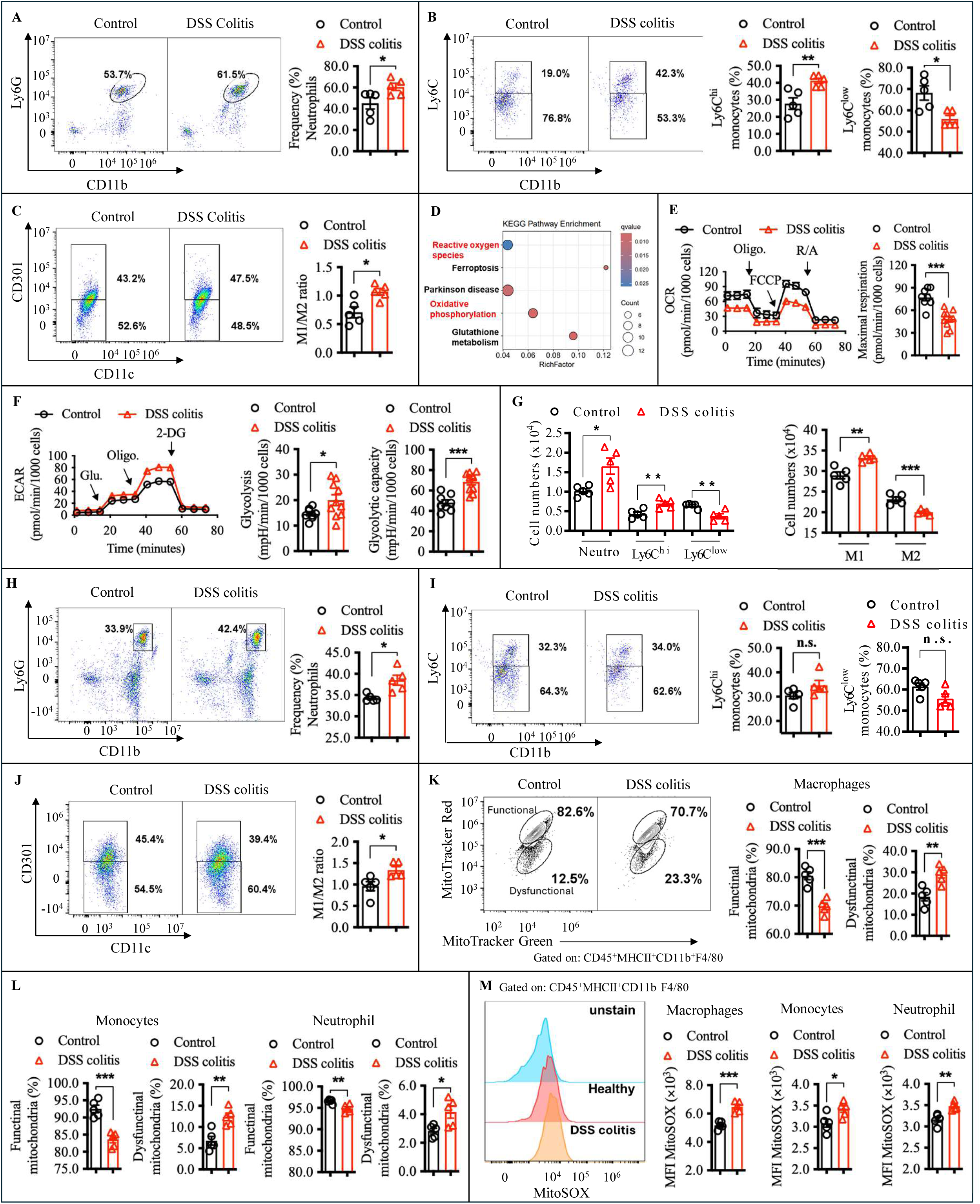
**Metabolic reprogramming of circulating immune cells in colitis. A-C**, Immune cells from blood were isolated on day 7 post-DSS and analyzed by flow cytometry. The percentage of neutrophils, Ly6C ^hi/low^ monocytes, and M1/M2 macrophages were quantified, and the M1/M2 ratio was calculated. n=5 mice/group. **D**, White blood immune cells were processed for RNA extraction and RNA-seq analysis. Pathway enrichment was performed using KEGG analysis. **E**&F, Oxygen consumption rate (OCR) and extracellular acidification rate (ECAR) of circulating immune cells were assessed using the Seahorse extracellular flux assay at day 7 post-DSS. **G–J**, Heart tissues collected at day 7 post-DSS were processed for flow cytometric analysis of innate immune cells, including quantification of absolute cell numbers (G) and percentages of neutrophils, monocytes, and macrophages (H-J). **K and L**, Heart tissues collected at day 42 post-DSS were analyzed by flow cytometry to assess mitochondrial function in heart-infiltrating immune cells. Mitochondrial mass was measured using MitoTracker Green, mitochondrial membrane potential using MitoTracker Red, and mitochondrial reactive oxygen species (mitoROS) production using MitoSOX Red. The proportions of cells with functional or dysfunctional mitochondria (K&L) and the mean fluorescence intensity (MFI) of mitoROS (L) were quantified in heart-infiltrating innate immune cells. n=5 mice/group. Data are presented as mean ± S.E.M. with statistical significance indicated as follows: n.s., not significant, *p*>0.05, * *p*<0.05, ** *p*<0.01, *** *p*<0.001.

We then examined whether these circulating immune alterations were accompanied by immune cell accumulation in the heart. At day 7 of colitis, flow cytometry showed increased number (Figure 2G) and percentages (Figure 2H-2J) of cardiac neutrophils, inflammatory monocytes and macrophages, with macrophages representing the predominant immune cell population in the heart (Figure 2G). By day 42, flow cytometry showed the percentage of cardiac macrophages, monocytes, and neutrophils remained elevated in colitis mice compared with controls, and H&E staining confirmed inflammatory cell infiltration in the heart (Figure S5E&S5F). Importantly, heart-infiltrating innate immune cells from colitis mice displayed marked mitochondrial dysfunction and elevated mitoROS production (Figure 2K-2M). Collectively, these findings demonstrate that colitis induce long-lasting mitochondrial dysfunction in systemic innate immune cells and promotes the accumulation of metabolically dysfunctional immune cells within cardiac tissue.

### Persistent Immune Cell Metabolic Dysfunction Drives Cardiac Impairment

Next, we investigated whether discontinuation of DSS treatment could resolve intestinal inflammation and reverse the systemic and cardiac abnormalities observed in colitis mice. DSS administration was halted, and mice were allowed a six-week recovery period with regular drinking water. Although the mice gradually regained body weight during this period (Figure 3A), histological and inflammatory marker analyses revealed persistent low-grade intestinal inflammation. This was characterized by continued immune cell infiltration in the lamina propria, crypt hyperplasia (Figure S6A), and elevated fecal lipocalin-2 levels (Figure S6B), indicating that subclinical intestinal inflammation persists despite apparent clinical recovery. In parallel with this residual intestinal inflammation, cardiac dysfunction also persisted. Echocardiography revealed reduced ejection fraction and fractional shortening (Figure 3B), Furthermore, morphological analyses demonstrated features of cardiac hypertrophy, including an increased heart weight and heart-to-body weight ratio (Figure 3C), and enlarged cardiomyocyte size (Figure 3D). Meanwhile, metabolic abnormalities in blood immune cells were also sustained after DSS withdrawal. Seahorse extracellular flux analysis revealed significantly reduced basal and maximal respiration (Figure 3E), accompanied by a compensatory increase in glycolysis and glycolytic capacity (Figure 3F). To examine the functional consequences of these metabolically reprogrammed immune cells on cardiomyocyte function and hypertrophy, we isolated circulating immune cells from control and DSS-treated mice and co-cultured them with H9C2 cardiomyocytes in a trans-well system, which permits paracrine signaling without direct cell contact (Figure 3G). Strikingly, H9C2 cells exposed to immune cells from colitis mice developed hypertrophic features, including increased cell surface area and protein content (Figure 3H and 3I), as well as elevated expression of hypertrophy-associated markers (ANP, BNP, and β-MHC) at both the mRNA and protein levels (Figure 3J and 3K).

**Figure 3.**
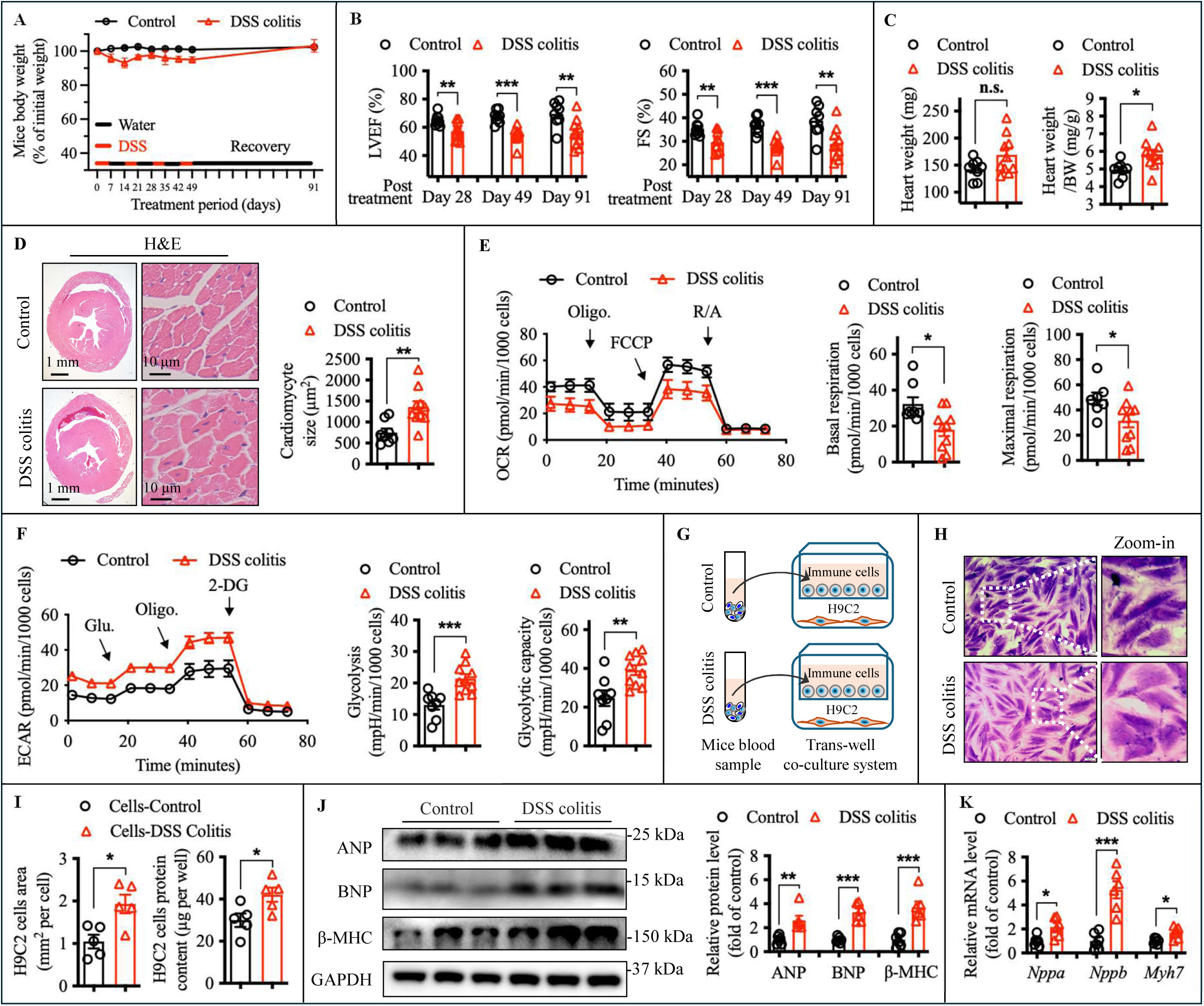
Incomplete recovery from colitis leads to persistent mitochondrial and cardiac dysfunction. DSS treatment was discontinued in colitis mice at week 7, followed by a 6-week recovery period during which mice received regular drinking water. **A**, Mice body weight throughout the experiment. n=8-10 mice/group. **B**, Cardiac function by ultrasound measurements of ejection fraction (LVEF) and fractional shortening (FS). n=8-10 mice/group. **C**, Actual mice heart weight and the ratio of heart weight to body weight (BW). n=8 mice/group. **D**, Representative images of H&E staining in the heart of mice, and quantification of cardiomyocyte size. n=8-10 mice/group. **E** and **F**, Oxygen consumption rate (OCR) and extracellular acidification rate (ECAR) of circulating immune cells were assessed using the Seahorse extracellular flux assay after 7 weeks of DSS cessation. n=8-10 mice/group. **G** through **K**, H9C2 cardiomyocytes were co-cultured with blood immune cells (**G**) isolated from control and colitis mice. Cardiomyocyte hypertrophy was assessed by measuring cell surface area, with representative images shown (**H**), and quantitative analysis of cell size and total protein content presented (**I**). Expression levels of hypertrophic markers ANP, BNP, and β-MHC were measured at the protein level by western blotting (**J**) and at the mRNA level by real-time PCR (**K**). Data are presented as mean ± S.E.M. with statistical significance indicated as follows: n.s., not significant, *p*>0.05, * *p*<0.05, ** *p*<0.01, *** *p*<0.001.

Since circulating immune cells derive from bone marrow hematopoietic stem and progenitor cells (HSPCs), we hypothesized that colitis impairs HSPC metabolism, thereby driving metabolic alterations in peripheral immune cells. Flow cytometry revealed reduced mitochondrial membrane potential in HSPCs (Figure S6C). Consistently, Seahorse analysis demonstrated decreased OCR, reduced maximal respiration, and increased ECAR, indicating colitis-induced mitochondrial dysfunction in HSPCs (Figure S6D and S6E). To determine whether these colitis-associated alterations in HSPCs contribute to metabolic dysfunction in peripheral immune cells and subsequent cardiac impairment, we performed bone marrow transplantation experiments. Lethally irradiated recipient mice were reconstituted with bone marrow from either colitis or healthy donors (Figure S6F). One and a half months later, mice receiving colitis donor marrow exhibited mitochondrial dysfunction in circulating immune cells, with reduced OCR and related metabolic parameters (Figure S6G and S6H), and developed cardiac dysfunction, including decreased LVEF, FS, LVIDd, and LVIDs (Figure S6I) as well as increased relative heart weight (Figure S6J). Together, these findings indicate that colitis induced a lasting metabolic imprint on immune cells, including bone marrow progenitors, that drive cardiac impairment.

### IBD Associated Gut Microbiota Alteration Contributes to Heart Dysfunction

Colitis is often linked to gut microbiota alterations that may contribute to cardiac dysfunction.^35,36^ To examine how DSS-induced colitis affects microbial composition, we performed 16S rRNA sequencing during active colitis (days 7 and 42) and recovery (days 91). β-Diversity analysis showed distinct clustering of control and DSS-treated groups (days 7 and 42), indicating microbial shifts; although, these differences did not reach statistical significance based on distance metrics (Figure S7A). α-Diversity measures, including Faith’s phylogenetic diversity and evenness, revealed a trend toward reduced species richness and a significant decrease in microbial evenness following DSS treatment (Figure S7B). Notably, despite clinical recovery after six weeks of DSS withdrawal, the microbiota remained altered at day 91, as evidenced by sustained β-diversity separation and incomplete restoration of microbial evenness compared with controls (Figure S7C). These alterations were further reflected in taxonomic analyses at the genus level. LEfSe analysis identified specific shifts in bacterial taxa, with significant enrichment of Thomasclavelia and depletion of Oscillibacter and Incertae Sedis in DSS-treated mice at day 7 post-colitis (Figure S7D). By day 42 and after six weeks of DSS withdrawal, additional taxa exhibited notable enrichment or depletion, indicating that DSS-induced colitis drives sustained remodeling of the gut microbial community (Figure S7E-S7F).

To test whether colitis-associated microbiota changes contribute causally to cardiac dysfunction, we performed fecal microbiota transplantation (FMT) from healthy or post-DSS colitis mice into germ-free (GF) recipients (Figure 4A). Two months post-FMT, mice receiving colitis-associated microbiota showed elevated levels of the intestinal inflammation marker fecal lipocalin-2 (LCN2), despite normal body weight, slightly reduced fat mass, and largely intact gut architecture (Figure S7G-S7I). These mice also exhibited systemic immune alterations, including increased circulating neutrophils and monocytes (Figure 4B), accompanied by reduced mitochondrial respiratory capacity in peripheral immune cells, including granulocytes and mononuclear cells (Figure 4C-4E). Accompanying these immune-metabolic changes, GF male mice receiving colitis-associated microbiota developed cardiac dysfunction, with reduced LVEF and FS (Figure 4G), as well as cardiac hypertrophy, evidenced by increased heart-to-body weight ratio, enlarged cardiomyocyte size, enhanced heart wall thickness, and elevated collagen volume (Figure 4F-4H). Collectively, these findings demonstrate that colitis-associated microbiota, even in the absence of overt clinical colitis, is sufficient to drive systemic immune-metabolic dysfunction and promote cardiac impairment.

**Figure 4.**
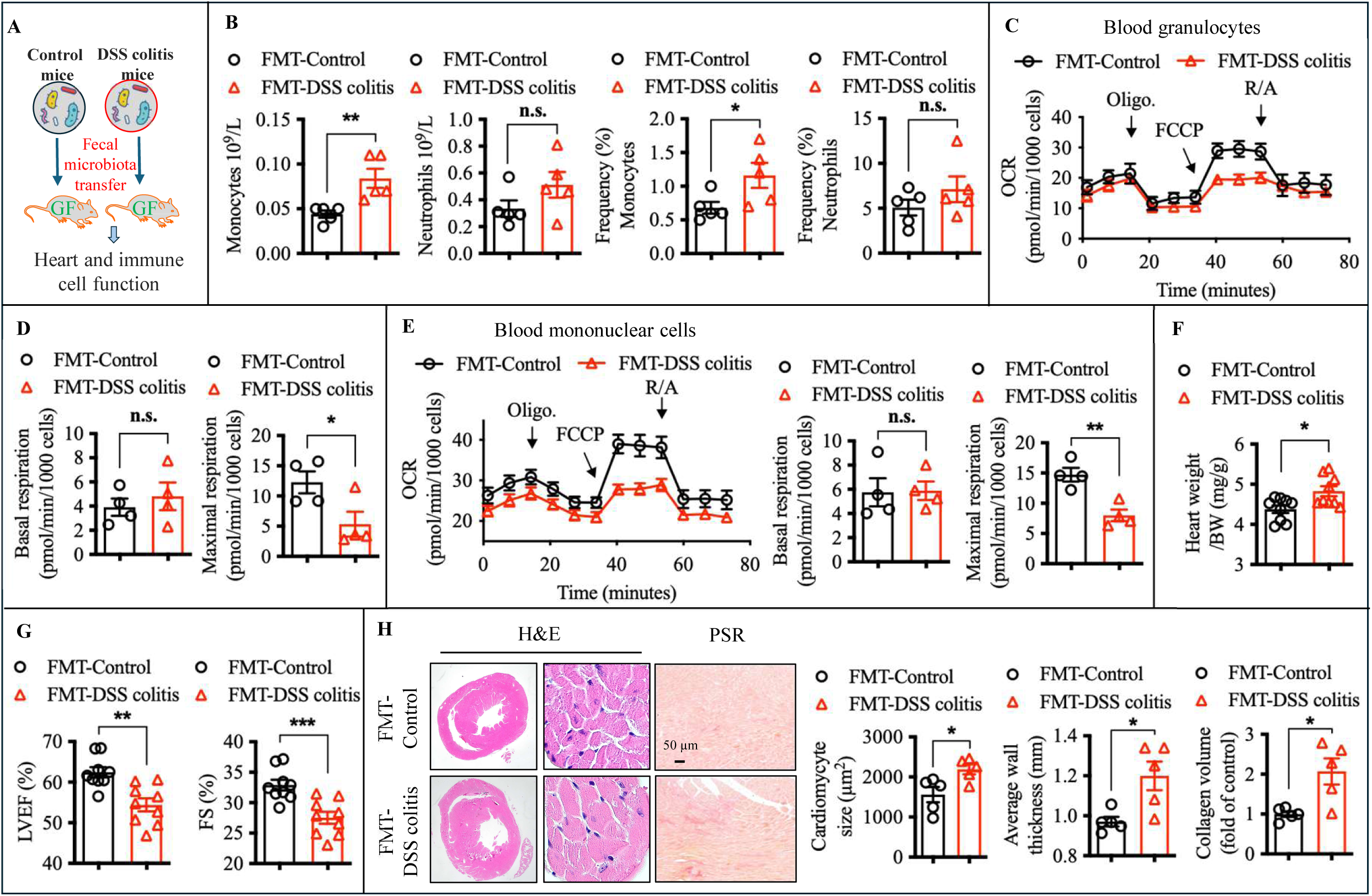
Colitis-associated gut microbiota induces immune and cardiac dysfunction in germ-free mice. **A** through **H**, Germ-free mice were administered a fecal microbiota transplant (FMT) from either healthy control or DSS colitis mice during the recovery phase at day 91. **A**, Scheme of the FMT experiment. **B**, Two months post-FMT, absolute numbers and percentage of neutrophils and monocytes in the blood were quantified using a hematology cell counter (n=5 mice/group). **C-E**, Blood granulocytes (**C** and **D**) and mononuclear cells (**E**) were isolated from FMT recipient mice using Ficoll-Paque Plus. Mitochondrial respiration, assessed by oxygen consumption rate (OCR), was measured using the Seahorse Analyzer (n=4 mice/group). **F**, The heart-to-body weight (BW) ratio (n=9 mice/group). **G**, Cardiac function in FMT recipient mice was evaluated by ultrasound, including measurements of ejection fraction (EF) and fractional shortening (FS). n=9 mice/group. **H**, Representative images of H&E and Picro Sirius Red (PSR) staining in the heart tissue, and quantification of cardiomyocyte size, average heart wall thickness, and collagen volume. n=5 mice/group. Data are presented as mean ± S.E.M. with statistical significance indicated as follows: n.s., not significant, *p*>0.05, * *p*<0.05, ** *p*<0.01, *** *p*<0.001.

### Colitis-Associated LPS Translocation Reprograms Immune Cell Metabolism *via* TLR4 to Drive Cardiac Dysfunction

To determine which innate immune cell populations drive colitis-associated cardiac dysfunction, we performed selective depletion experiments. Neutrophils were depleted using a well-characterized anti-Ly6G antibody, whereas macrophages were depleted using clodronate liposomes and anti-CSF1R treatment, as shown in Figure S8A. Notably, neutrophil depletion did not prevent colitis-induced cardiac dysfunction, whereas macrophage depletion markedly ameliorated cardiac dysfunction (Figure S8B). We next sought to identify circulating factors, particularly microbiota-derived components, that mediate macrophage activation. To this end, we performed functional screening by treating macrophages with serum from control or DSS-treated mice. To define the biochemical nature of the active component(s), we subjected serum to selective fractionation (Fig S8C). Notably, heat inactivation, Proteinase K digestion, and DNase/RNase treatment failed to diminish the pro-inflammatory activity of DSS serum, arguing against protein- or nucleic acid-mediated effects (Fig S8D). In contrast, lipid removal markedly reduced this activity, suggesting involvement of a lipid-containing, heat-stable, non-proteinaceous factor, compatible microbiota-derived endotoxins such as LPS (Fig S8D). Supporting this interpretation, addition of the LPS-neutralizing agent polymyxin B markedly attenuated TNF-α induction in macrophages (Figure S8E), implicating LPS as a major contributor.

Supporting this, circulating LPS levels were significantly elevated in DSS-treated mice (Figure 5A). Given that macrophages underlie chronic colitis-induced cardiac dysfunction and that circulating LPS is elevated, we next asked whether LPS signals through hematopoietic Toll-like receptor 4 (TLR4) to drive immune cell reprogramming and cardiac pathology. To address this, we generated bone marrow chimeras by reconstituting irradiated wild-type (WT) recipients with WT (BM-WT) or TLR4-deficient (BM-TLR4 KO) bone marrow (Figure 5B). In both groups, non-hematopoietic cells, including cardiomyocytes, remain WT. In the BM-WT group, hematopoietic cells express TLR4 and serve as controls, whereas in the BM-TLR4 KO group, hematopoietic cells specifically lack TLR4. Following DSS-induced colitis and recovery, BM-WT and BM-TLR4KO colitis mice showed systemic metabolic parameters comparable to their respective non-colitis controls, including blood glucose and cholesterol levels, with lower triglyceride levels observed in colitis mice (Figure S9A). However, circulating immune cells from BM-WT colitis mice exhibited metabolic reprogramming compared with BM-WT non-colitis controls, characterized by a modest reduced oxygen consumption rate (OCR), increased extracellular acidification rate (ECAR), and broad bioenergetic alterations (Figure 5C-5F). In contrast, circulating immune cells from BM-TLR4 KO colitis mice maintained near-normal metabolic profiles largely comparable to BM-TLR4 KO non-colitis controls (Figure 5C-5F). Along with these metabolic changes, circulating immune cells from WT colitis mice induced robust hypertrophy when co-cultured with H9C2 cardiomyoblasts, whereas immune cells from TLR4-deficient colitis mice failed to elicit this response (Figure 5G). In vivo, BM-WT colitis mice developed significant cardiac dysfunction, as indicated by reduced LVEF and FS, increased LVIDd and LVIDs, elevated relative heart weight, cardiomyocyte hypertrophy, increased ventricular wall thickness, and augmented collagen deposition compared with BM-WT non-colitis controls (Figure 5H-5K). In contrast, BM-TLR4 KO colitis mice displayed minimal cardiac impairment and structural alterations compared with BM-TLR4 KO non-colitis controls (Figure 5H-5K), further supporting a critical role for hematopoietic TLR4 signaling in linking colitis-associated immune cells alteration to cardiac dysfunction.

**Figure 5.**
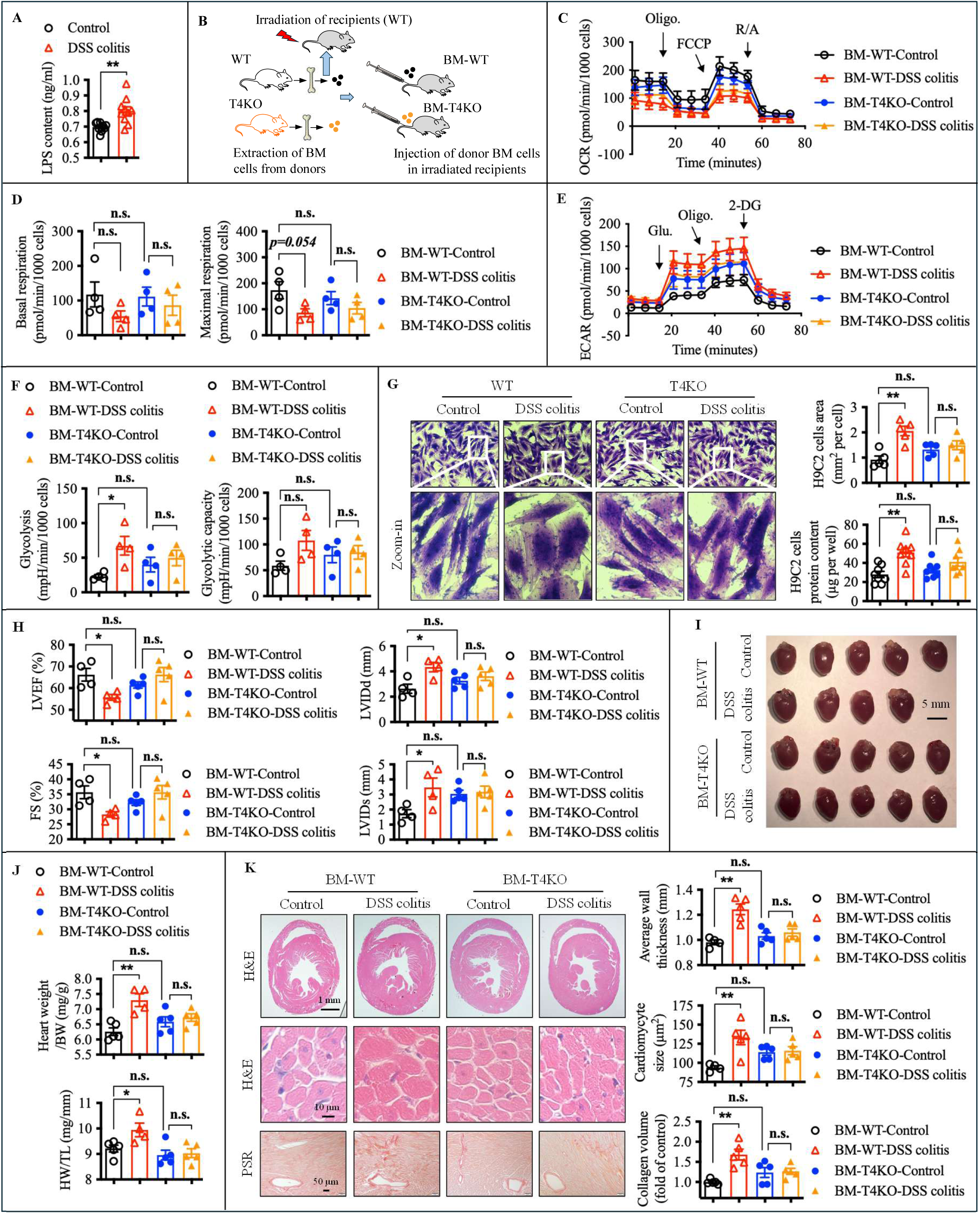
Gut bacteria-derived LPS reprograms immune cell metabolism through TLR4 signaling to impair cardiac function. **A**, Circulating lipopolysaccharide (LPS) levels were measured in DSS-treated mice at day 7 post DSS treatment. n=10 mice/group. **B**, Schematic of bone marrow chimeric mice generated by irradiating WT recipients and reconstituting them with either WT (BM-WT) or TLR4 KO (BM-T4KO) bone marrow to restrict TLR4 deficiency to hematopoietic cells. **C** through **K**, BM-WT and BM-T4MKO mice were treated with DSS for four cycles to induce colitis, followed by a 6-week recovery phase. **C** through **F**, Circulating immune cells were isolated and analyzed for oxygen consumption rate (OCR) and extracellular acidification rate (ECAR) using Seahorse extracellular flux assays. **G**, H9C2 cardiomyocytes were co-cultured with blood immune cells isolated from WT and TLR4 KO mice (T4KO), with or without colitis. Cardiomyocyte hypertrophy was assessed by measuring cell surface area, with representative images shown and quantitative analysis of cell size and total protein content presented (n=5). **H**, Cardiac function was assessed by echocardiography measuring ejection fraction (EF), fractional shortening (FS), LV end-diastolic internal dimension (LVIDd), and LV end-systolic internal dimension (LVIDs). n=4-5 mice/group. **I** and **J**, The ratios of heart weight (HW) to body weight and heart weight to tibia length (TL). n=4-5 mice/group. **K**, Representative images of H&E staining in the heart tissue, with quantification of average ventricular wall thickness and cardiomyocyte size, alongside Picro Sirius Red (PSR) staining. n=4-5 mice/group. Data are presented as mean ± S.E.M. with statistical significance indicated as follows: n.s., not significant, *p*>0.05, * *p*<0.05, ** *p*<0.01, *** *p*<0.001.

IBD is frequently associated with an increased abundance of *Escherichia coli* strains enriched in lipopolysaccharide (LPS).^37^ To determine whether colonization with such bacteria is sufficient to induce immune and cardiac alterations via hematopoietic TLR4 signaling, we colonized BM-WT and BM-TLR4 KO mice with an LPS-positive (LPS⁺) pathobiont *E. coli* strain. In BM-WT mice, colonization induced immune metabolic reprogramming, characterized by reduced oxygen consumption rate (OCR) and increased extracellular acidification rate (ECAR), and was accompanied by moderate cardiac dysfunction (Fig. S9B-S9E). In contrast, BM–TLR4 KO mice were protected from these metabolic and cardiac alterations (Fig. S9B-S9E). Moreover, continuous systemic LPS infusion via subcutaneous osmotic minipump impaired cardiac function, further demonstrating that sustained LPS exposure alone is sufficient to drive cardiac dysfunction (Fig. S9F-S9J). Together, these findings demonstrate that colitis-associated LPS translocation reprograms circulating immune cell metabolism through hematopoietic TLR4 signaling, ultimately leading to cardiomyocyte hypertrophy and cardiac dysfunction.

### GBP1 Mediates LPS-Induced Mitochondrial Fission by Promoting DRP1 Mitochondrial Recruitment

Next, we sought to investigate how LPS influences metabolic alterations in innate immune cells that may contribute to cardiac dysfunction. Based on our RNA-seq analysis of circulating immune cells from control and colitis mice, GBP1 emerged as one of the most differentially expressed genes (Figure 6A). Upregulation of GBP1 was further confirmed in circulating immune cells by qRT-PCR and ELISA (Figure 6B and 6C), whereas other GBPs were comparable between groups (Figure S10A). Immunohistochemistry further revealed GBP1 expression in heart tissue, particularly within cardiac macrophages of colitis mice (Figure S10B and S10C). GBP1 upregulation in circulating immune cells from WT colitis mice was abolished in TLR4 KO colitis mice (Figure 6D), indicating LPS-TLR4-dependent induction. *In vitro*, LPS treatment of THP-1 monocytes, even at low doses (10 ng/mL), robustly induced GBP1 expression within 4 hours, whereas untreated cells showed no detectable GBP1 (Figure 6E-6G). To investigate whether and how GBP1 contributes to mitochondrial and metabolic changes in immune cells, we first accessed its subcellular localization during LPS stimulation. Western blot analysis of whole-cell, cytosolic, and mitochondrial fractions revealed predominant localization of GBP1 to mitochondria after LPS treatment (Figure S10D). Immunoprecipitation and confocal microscopy further confirmed that GBP1 physically associates and colocalizes with dynamin-related protein 1 (DRP1), a cytosolic GTPase that translocate to mitochondria to mediate fission, at mitochondria (Figure 6H through 6J). GBP1 also colocalized with fission 1 protein (FIS1), the mitochondrial receptor that anchors DRP1 at fission sites (Figure S10E). Functionally, LPS-induced DRP1 recruitment to mitochondria was abolished in GBP1-deficient THP-1 cells, without affecting total DRP1 levels (Figure 6K-6N), demonstrating that GBP1 is required for DRP1 mitochondrial translocation. Consistently, LPS treatment caused pronounced mitochondrial fragmentation, reflected by reduced mitochondrial length and increased fission, which was largely prevented by GBP1 knockdown (Figure 6O through 6Q). Together, these results demonstrate that GBP1 links LPS signaling to DRP1 mitochondrial recruitment and mitochondrial fission, thereby promoting mitochondrial morphological remodeling in immune cells.

**Figure 6.**
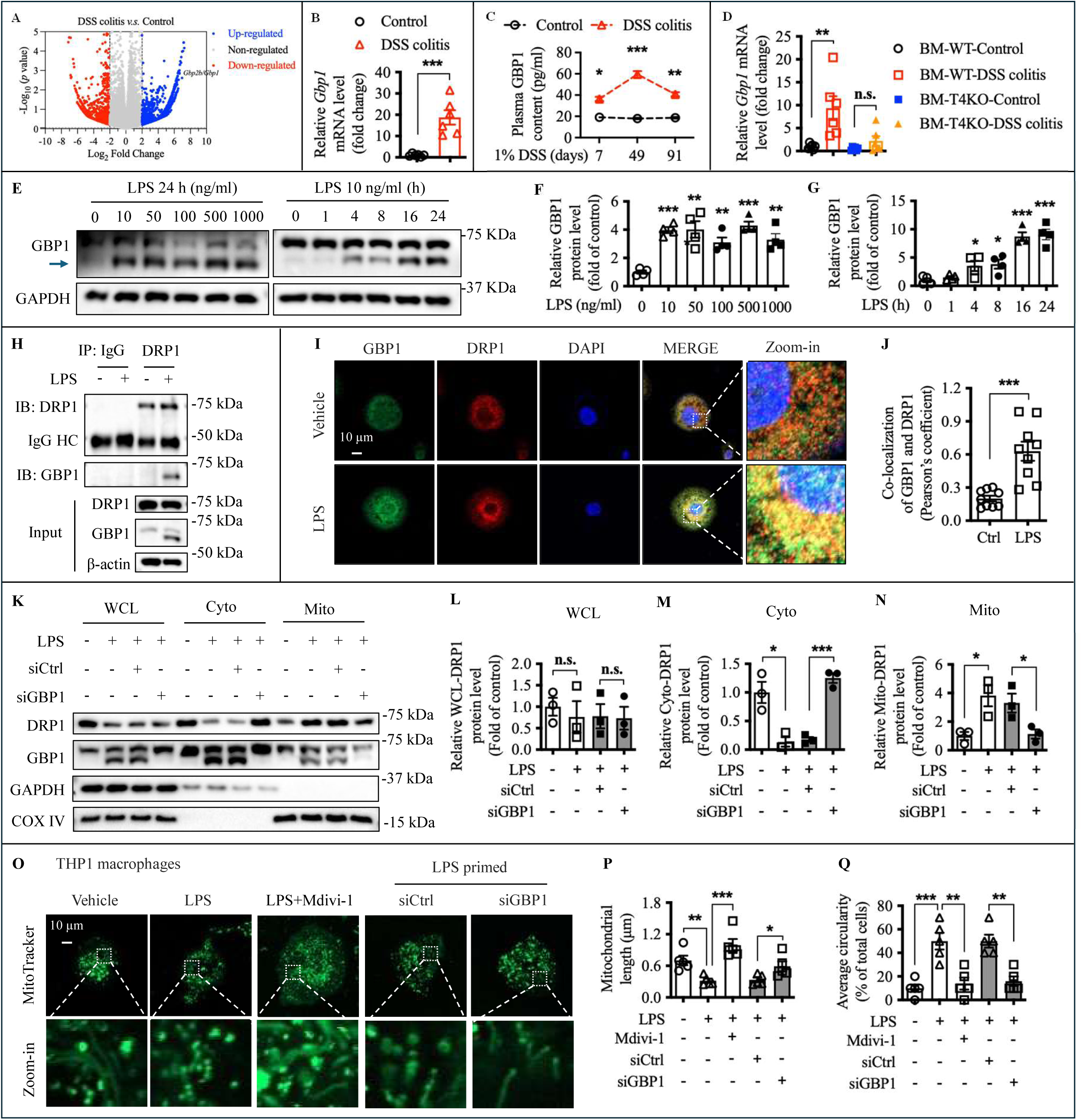
GBP1 mediates LPS-induced DRP1-dependent mitochondrial fission. **A**, RNA sequencing (RNA-seq) analysis of circulating immune cells from mice treated with DSS for 7 days, with differentially expressed genes visualized by a volcano plot. **B**, Quantitative RT PCR analysis of guanylate-binding protein 1 (GBP1) expression in circulating immune cells from mice treated with DSS for 7 days. **C**, Plasma GBP1 levels in control and DSS-induced colitis mice were measured by ELISA at days 7, 49, and 91 after DSS treatment. **D**, GBP1 expression in circulating immune cells from BM-WT and BM-TLR4 KO colitis mice. **E** through **G**, THP1 macrophages were treated with indicated concentrations (0, 10, 50, 100, 500, and 1000 ng/ml) LPS for 24h or treated with 10 ng/ml LPS for indicated time intervals (0, 1, 4, 8, 16, and 24 h). Then GBP1 protein expression was detected by western blot analysis. n=4. **H** through **J**, THP1 macrophages were treated with or without 10 ng/ml LPS for 24h. The interaction between GBP1 and DRP1 were then evaluated by immunoprecipitation (IP) analysis using whole- cell lysates (**H**) and by immunofluorescence (IF) staining (**I** and **J**, n=9). **K** through **N**, THP1 macrophages were primed with 10 ng/ml LPS for 8 h and were then transfected with control siRNA (siCtrl) or GBP1 siRNA (siGBP1) *via* electroporation. After 48h, cell fractional analysis was performed to detect the protein levels of DRP1 and GBP1 in the fractions of whole cell lysate (WCL, **L**), cytoplasm (Cyto, **M**), and mitochondria (Mito, **N**), respectively. n=3. **O** through **Q**, THP1 macrophages were primed with 10 ng/ml LPS for 8 h and were then transfected with control siRNA (siCtrl) or GBP1 siRNA (siGBP1) *via* electroporation. Mitochondrial Division inhibitor 1 (Mdivi-1, 50 µM) was used as positive control. After 48h, mitochondrial morphology was viewed by staining of Mitotracker green (**O**), the mitochondrial length (**P**) and average circularity percentage (**Q**) were quantified. n= 5. Data are presented as mean ± S.E.M. with statistical significance indicated as follows: n.s., not significant, *p*>0.05, * *p*<0.05, ** *p*<0.01, *** *p*<0.001.

### GBP1 Drives LPS-Induced Mitochondrial Dysfunction and Macrophage Recruitment to The Heart

Given the close link between mitochondrial fission, metabolic function, and reactive oxygen species (ROS) production,^38^ and our finding that GBP1 mediates LPS-induced mitochondrial fission, we next investigated whether GBP1 also contributes to LPS-induced metabolic dysfunction and ROS generation. Seahorse analysis showed that GBP1 knockdown during LPS stimulation restored both maximal respiration and glycolytic capacity, similar to treatment with Mdivi-1, a Drp1 inhibitor that prevents mitochondrial fission (Figure 7A-7C). Consistently, LPS treatment significantly increased MitoROS in THP-1 cells, whereas GBP1 knockdown largely prevented this elevation, indicating that GBP1 is required for LPS-induced mitochondrial oxidative stress (Figure 7D). Because mitochondrial dysfunction profoundly influences macrophage behavior, we next assessed whether GBP1 also regulates immune cell migration. GBP1 knockdown significantly reduced THP-1 migration during LPS stimulation *ex vivo*, as shown by Trans-well and scratch assays (Figure 7E -7G). To determine whether GBP1 similarly regulates macrophage recruitment to the heart *in vivo*, we adoptively transferred CD45.1⁺ bone marrow-derived macrophages, with or without GBP1 deletion, into CD45.2 recipient mice intraperitoneally, followed by systemic LPS administration. Twelve hours after LPS injection, hearts were analyzed for the presence of transferred CD45.1⁺ macrophages (Figure 7H). GBP1-deficient macrophages exhibited markedly reduced infiltration into the heart, reflected by decreased cell numbers and frequencies, and this impaired recruitment was associated with reduced expression of the chemokine receptor CCR2 (Figure 7I-7L). Together, these results identify GBP1 as a critical mediator of LPS-induced mitochondrial dysfunction, oxidative stress, and macrophage infiltration into the heart.

**Figure 7.**
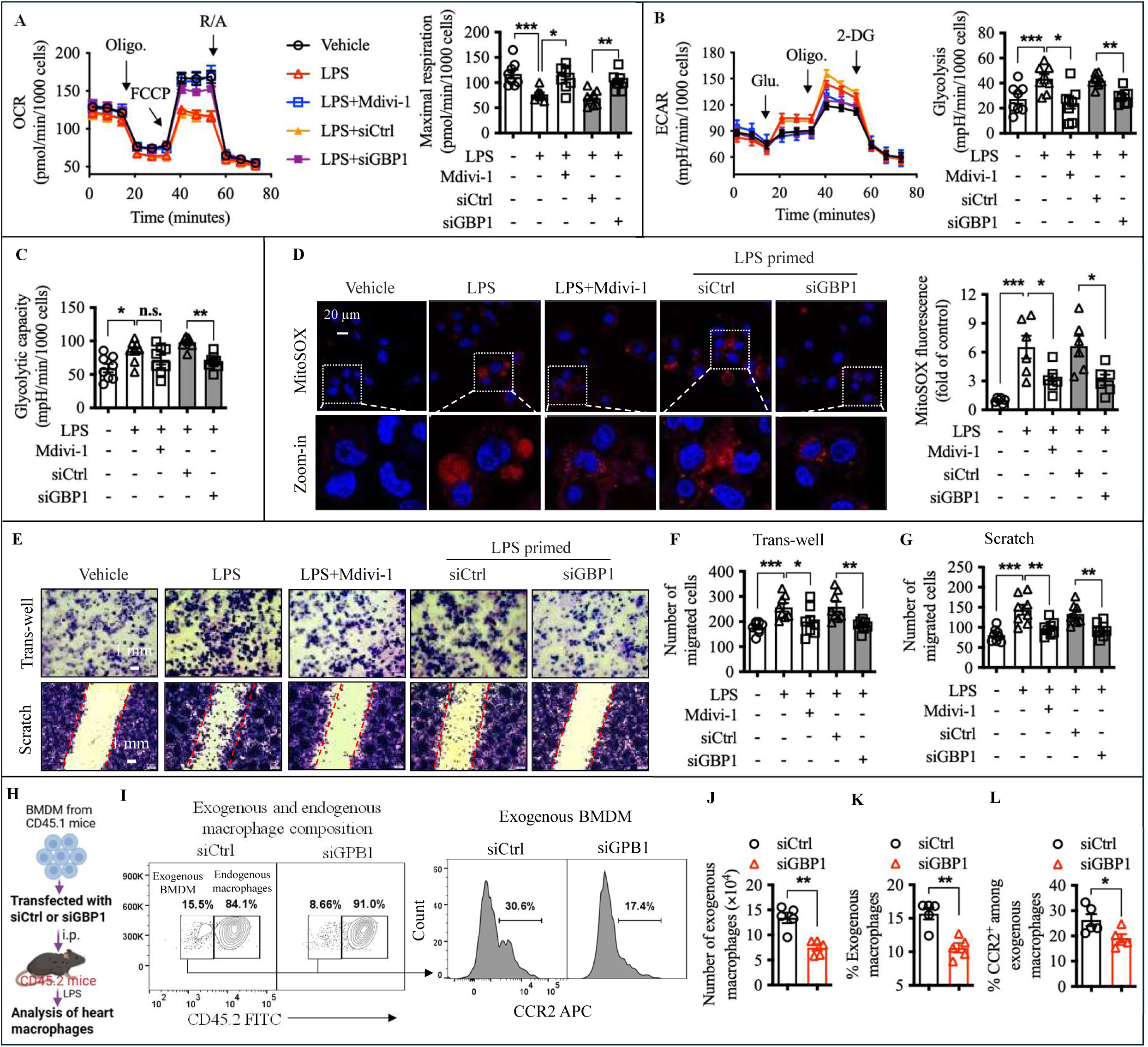
GBP1 is required for LPS-induced macrophage metabolic reprogramming and infiltration. **A** through **G**, THP1 macrophages were primed with 10 ng/ml LPS for 8 h and were then transfected with control siRNA (siCtrl) or GBP1 siRNA (siGBP1) *via* electroporation. Mitochondrial Division inhibitor 1 (Mdivi-1, 50 µM) was used as positive control. **A** through **C**, Oxygen consumption rate (OCR) and extracellular acidification rate (ECAR) were measured using Seahorse extracellular flux assays (n=8). **D**, MitoSOX level was detected by IF staining (n=6). **E** through **G**, Trans-well invasion assay and scratch wound healing assay were performed and the number of migrated cells were quantified (n=8). **H** through **L**, CD45.1 bone marrow-derived macrophages (BMDM) were transfected with control siRNA (siCtrl) or GBP1 siRNA (siGBP1) *via* electroporation and were then injected into CD45.2 recipient mice prior to LPS administration. Exogenous and endogenous macrophage composition were determined by flow cytometry. n= 5 mice/group. Data are presented as mean ± S.E.M. with statistical significance indicated as follows: n.s., not significant, *p*>0.05, * *p*<0.05, ** *p*<0.01, *** *p*<0.001.

### Exosomal GBP1 Mediates Colitis/Inflammation-Induced Cardiac Dysfunction and Hypertrophy

Exosomes are small extracellular vesicles that transfer bioactive molecules between cells and thereby influence cardiac remodeling, and hypertrophy, particularly under chronic inflammatory conditions.^39^ We next isolated exosomes from the serum of colitis mice and found that they contained elevated levels of GBP1 (Figure 8A). To determine whether circulating exosomes can distribute systemically and reach the heart, thereby potentially delivering GBP1, we performed in vivo tracking and uptake analyses using DiR labeling. Whole-body IVIS imaging at 2 h and 24 h post-injection showed that free DiR dye was predominantly retained in the liver, consistent with rapid hepatic clearance (Figure S11A). In contrast, DiR-labeled exosomes displayed a broader distribution pattern, with detectable signals across multiple anatomical regions (Figure S11A). Ex vivo imaging further confirmed that exosomes accumulated in several organs, including the liver, spleen, lung, and notably the heart (Figure S11B). Consistent with these findings, fluorescence imaging of tissue sections revealed DiR-positive signals in heart tissue following exosome administration, whereas minimal signal was observed with free dye controls (Figure S11C). Moreover, in vitro assays demonstrated that DiR-labeled exosomes were efficiently internalized by H9C2 cardiomyocytes, as evidenced by robust intracellular fluorescence after 24 h incubation (Figure S11D). Together, these results show that circulating exosomes reach and are internalized by cardiomyocytes, enabling delivery of bioactive cargo such as GBP1.

**Figure 8.**
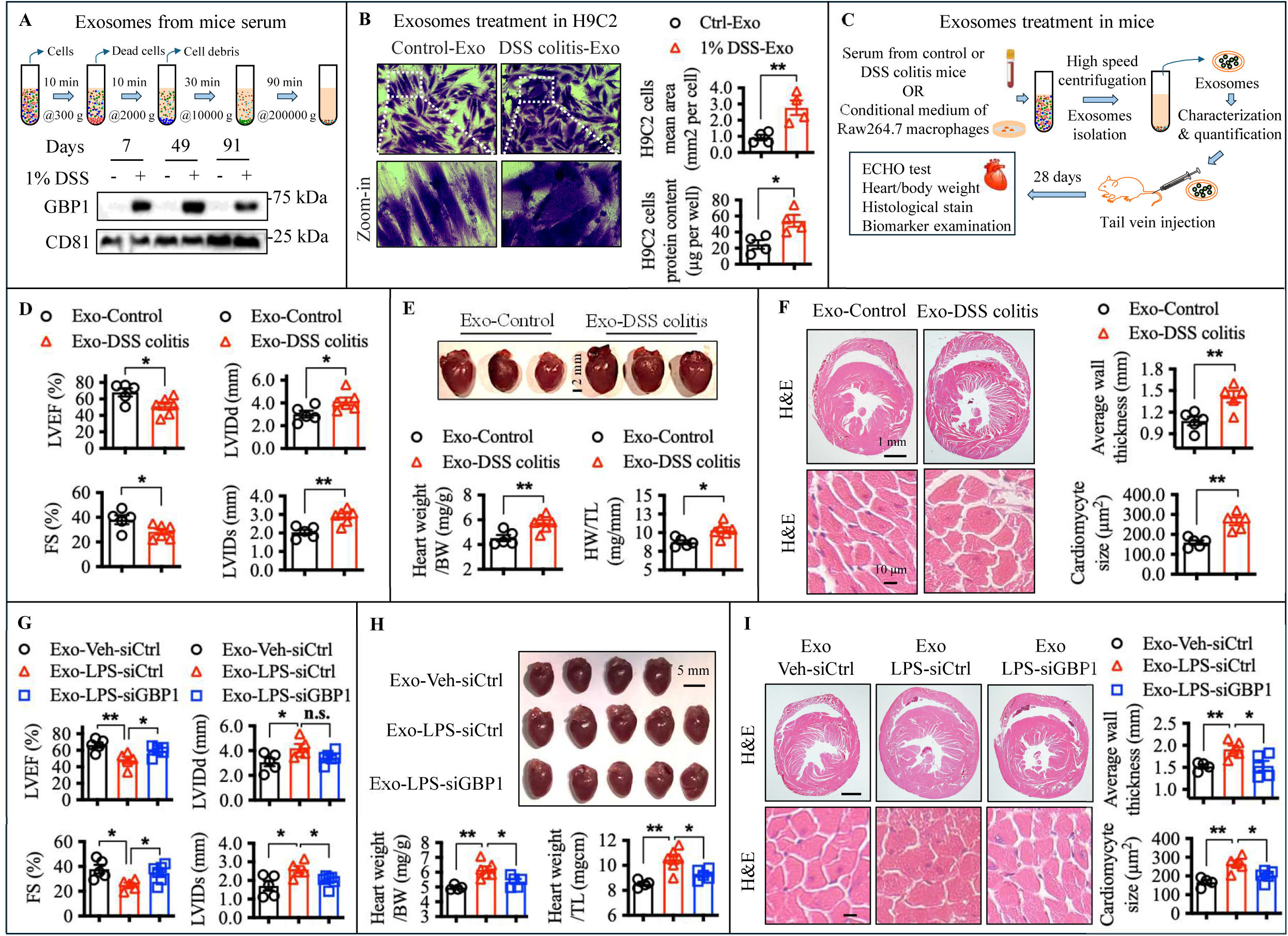
Exosomal GBP1 mediates inflammation-induced cardiac dysfunction and hypertrophy. **A**, Exosomes were isolated from the serum of mice in different stages of 1% DSS-induced colitis. Exosomal GBP1 was determined by western blot. CD81 was used as exosome marker. **B**, H9C2 cardiomyocytes were co-cultured with exosomes isolated from the serum of control or DSS colitis mice. Cardiomyocyte hypertrophy was assessed by measuring cell surface area with representative images shown and quantitative analysis of cell size and total protein content presented (n=4). **C**, Scheme of exosome treatment in mice. **D** through **F**, WT mice were treated with exosomes isolated from serum of control or DSS colitis mice *via* tail vein injection (30 µg/mouse/3 days) for 28 days. n=5-6 mice/group. **D**, Cardiac function was evaluated by measurements of ejection fraction (EF) and fractional shortening (FS). **E**, Representative images of whole heart and quantification of the ratios of heart weight (HW) to body weight (BW) and HW to tibia length (TL). **F**, Representative images of H&E staining in the heart tissue, and quantification of cardiomyocyte size as well as the average heart wall thickness. **G** through **I**, WT mice were treated with exosomes isolated from conditional medium of control siRNA (siCtrl)- or GBP1 siRNA (siGBP1)-transfected Raw264.7 macrophages (primed by 10 ng/ml LPS for 8 h) *via* tail vein injection (30 µg/mouse/3 days) for 28 days. Exosomes isolated from vehicle-treated and siCtrl-transfected macrophages was used as control. n=5-6 mice/group. **G**, Cardiac function was evaluated by measurements of ejection fraction (EF) and fractional shortening (FS). **H**, Representative images of whole heart and quantification of the ratios of heart weight (HW) to body weight (BW) and HW to tibia length (TL). **I**, Representative images of H&E staining in the heart tissue, and quantification of cardiomyocyte size as well as the average heart wall thickness. Data are presented as mean ± S.E.M. with statistical significance indicated as follows: n.s., not significant, *p*>0.05, * *p*<0.05, ** *p*<0.01.

To assess the functional consequences of this exosome-mediated signaling, co-culture of H9C2 cardiomyocytes with these serum-derived exosomes was sufficient to induce cardiomyocyte hypertrophy compared with exosomes from healthy control mice (Figure 8B). Moreover, systemic administration of exosomes from colitis mice into healthy recipients triggered cardiac dysfunction, evidenced by reduced LVEF FS, accompanied by overt cardiac hypertrophy (Figure 8C-8F; Figure S12A). To define the specific contribution of GBP1 to exosome-mediated cardiomyocyte remodeling, we stimulated human THP-1 cells with LPS and found that GBP1 was released into the culture medium *via* exosomes, whereas GBP1 silencing abolished its exosomal presence (Figure S12B). Treatment of H9C2 cardiomyocytes with GBP1-containing exosomes induced robust hypertrophic responses, reflected by increased cell size and total protein content. In contrast, exosomes derived from GBP1-silenced THP-1 cells failed to elicit hypertrophy (Figure S12C-S12E), demonstrating that exosomal GBP1 is required for cardiomyocyte hypertrophic signaling. To evaluate this effect *in vivo*, mice were administered exosomes containing GBP1 or GBP1-deficient exosomes from Raw264.7 macrophages. Mice receiving GBP1-containing exosomes developed marked cardiac hypertrophy and functional decline, whereas those treated with GBP1-deficient exosomes showed minimal hypertrophy and preserved cardiac function (Figure 8G-8I; Figure S12F), confirming that exosomal GBP1 is a key driver of colitis- and inflammation-associated cardiac remodeling. Together, these results demonstrate that GBP1 is secreted *via* exosomes and functions as a critical mediator of cardiac remodeling during inflammatory conditions.

## DISCUSSION

While epidemiological studies have established that patients with IBD are at increased risk for cardiovascular complications, the mechanisms linking chronic intestinal inflammation to cardiac injury remain incompletely understood. Our findings in a chronic colitis mouse model recapitulate the clinical link between intestinal inflammation and cardiac dysfunction, demonstrating that colitis impairs cardiac function in a gut microbiota-dependent manner. Colitis-associated dysbiosis leads to translocation of LPS, which drives metabolic reprogramming of innate immune cells, particularly macrophages, enhancing their migratory potential and facilitating cardiac infiltration. Importantly, we identified GBP1 colocalized with DRP1 in mitochondria, acting as a key mediator of LPS-induced mitochondrial fragmentation, metabolic reprogramming, and immune cell migration. Notably, GBP1 is released in serum exosomes during colitis, through which it is delivered to cardiomyocyte and induced hypertrophy, serving as a second hit (Figure 9). These results highlight the LPS-TLR4-GBP1 axis as a central pathway mediating colitis-associated cardiovascular risk and a potential therapeutic target.

**Figure 9.**
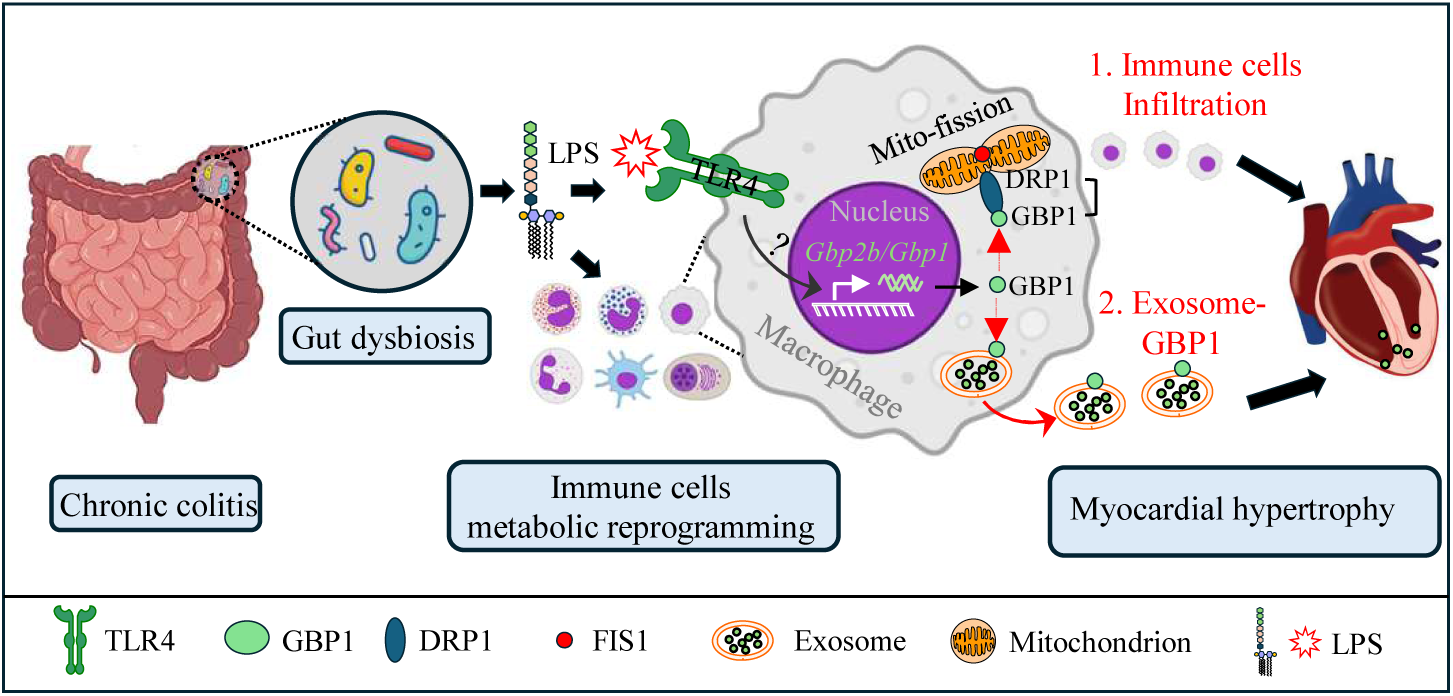
Proposed mechanism by which chronic colitis promotes myocardial hypertrophy. Colitis-associated dysbiosis leads to translocation of LPS, which induces GBP1 expression in immune cells. GBP1 colocalizes with DRP1 in mitochondria, driving mitochondrial fragmentation and metabolic reprogramming, thereby enhancing immune cell migratory capacity and promoting cardiac infiltration. Concurrently, GBP1 is released in serum exosomes during colitis, through which it is delivered to cardiomyocyte and induces hypertrophy, acting as a second hit.

In IBD, including ulcerative colitis and Crohn’s disease, the disease course typically follows a relapsing-remitting pattern.^40,41^ Although symptoms may improve or resolve during remission, underlying biological abnormalities often persist.^42,43^ Consistent with this, our chronic colitis mouse model revealed that colitis-associated alterations in the gut microbiota persist even after clinical remission. Fecal microbiota transplantation into GF mice further showed that these dysbiosis microbial communities are sufficient not only to induce intestinal inflammation but also to impair cardiac function, indicating that persistent microbial alterations may act as a sustained driver of both intestinal and extraintestinal pathology. Moreover, we demonstrated that *E. coli*, a genus frequently enriched in patients with IBD,^44,45^ can induce cardiac dysfunction when transferred into mice, supporting a direct mechanistic link between specific dysbiosis taxa and cardiovascular injury. In addition, our bone marrow transplantation experiments revealed that colitis or its associated gut microbiota impose a lasting metabolic imprint on bone marrow progenitors, which may also contribute to the persistently elevated cardiovascular risk observed even during periods of clinical remission.

The gut microbiome is increasingly recognized as a contributor to CVD, largely through microbial metabolites and structural components that translocate into the circulation and influence host physiology.^46,47^ Epidemiological studies show that elevated circulating LPS or LPS-binding protein, a stable proxy for endotoxin exposure, is associated with increased risk of atherosclerosis, myocardial infarction, and cardiac dysfunction, highlighting the clinical relevance of gut-derived endotoxemia-TLR4 signaling in humans.^48^ Consistent with this, prior studies have implicated TLR4 signaling in pathological cardiac remodeling, including angiotensin II-induced inflammatory remodeling and cardiac hypertrophy through TLR4/TRIF-dependent pathways^49,50^, as well as neonatal hyperoxia-induced left ventricular hypertrophy and dysfunction that can be prevented by systemic TLR4 antagonism.^51^ However, the cellular source through which TLR4 drives cardiac dysfunction may be context-dependent. For example, previous studies have shown that LPS can directly activate TLR4 signaling in endothelial cells or cardiomyocytes to promote vascular and cardiac inflammation.^52,53^ In contrast, our findings suggest that the modest but chronic elevation of LPS observed during colitis impairs cardiac function primarily through immune cell-mediated mechanisms. This context dependent effect may reflect differential cellular sensitivity, whereby high concentrations of LPS directly affect cardiac cells, whereas lower chronic elevations preferentially activate sensitive immune cell populations that secondarily promote cardiac dysfunction.

Peripheral metabolic organs can influence heart function through systemic metabolic and inflammatory pathways. DSS-induced colitis has been reported to exacerbate hepatic inflammation and fibrogenesis under certain metabolic disease conditions, such as experimental NASH, and liver metabolic dysfunction can contribute to cardiac remodeling and hypertrophy. ^54^ Other metabolic organs may also affect cardiac function by regulating lipid metabolism, glucose homeostasis, inflammatory mediators, and systemic energy balance.^55,56^ However, in our chow-fed DSS colitis model, we did not observe evidence of overt metabolic organ injury or systemic metabolic dysfunction. Instead, our data supports a model in which gut barrier disruption and low-grade endotoxemia activate hematopoietic immune cells to promote cardiac inflammation and dysfunction. Consistent with this immune cell-centric mechanism, circulating immune cells from DSS-treated mice showed marked bioenergetic dysfunction, with maximal respiration reduced below basal respiration and therefore a negative spare respiratory capacity. This likely reflects severe mitochondrial impairment, in which chronic inflammatory and endotoxin exposure exhausts mitochondrial reserve and blunts the response to FCCP-mediated uncoupling, a pattern consistent with prior reports that reduced spare respiratory capacity reflects impaired mitochondrial adaptation under metabolic stress.^57,58^

Evidence indicates that immune cells and their secreted cytokines play central roles in the development and progression of CVD.^59^ Proinflammatory cytokines, including IL-1β, IL-6, and interferons, are known to promote endothelial dysfunction, vascular inflammation, and adverse cardiac remodeling.^60,61^ Our study further identifies the cytokine-inducible GTPase GBP1 as a critical mediator linking chronic gut inflammation to cardiac dysfunction. While GBP1 is well-established for its antimicrobial functions, including destabilization of pathogen-containing vacuoles and promotion of inflammasome activation,^18,62^ we uncover a novel role for GBP1 in mediating LPS-induced mitochondrial fission and immunometabolism reprogramming in immune cells *via* DRP1-dependent mitochondrial translocation. Although the precise molecular details of the GBP1-DRP1 interaction remain to be fully elucidated, this pathway represents a previously unrecognized mechanism by which GBP1 drives mitochondrial dysfunction, enhances immune cell migratory potential, and contributes to cardiac dysfunction.

Beyond its intracellular role in driving immune cell metabolic reprogramming, GBP1 was also found to be secreted in exosomes, which directly induce cardiomyocyte hypertrophy. Exosomes are key mediators of cardiac hypertrophy, carrying microRNAs, proteins, and lipids that regulate cardiomyocyte growth, fibrosis, and inflammation.^39,63^ Despite challenges related to tissue targeting and cargo heterogeneity, exosomes remain both critical contributors to cardiac remodeling and attractive therapeutic targets in hypertrophic heart disease.^64^ In our study, we demonstrated that colitis or LPS stimulation leads to GBP1 incorporation into exosomes, which in turn promote cardiomyocyte hypertrophy both *in vitro* and *in vivo*. While prior studies have largely focused on soluble immune cell-derived cytokines as mediators of cardiac hypertrophy,^65,66^ our data reveal that cytokine-induced GBP1 packaged within exosomes can directly promote cardiomyocyte hypertrophy and functional impairment. This exosome-based signaling mechanism may confer greater stability, tissue reach, thereby enabling sustained cardiac effects even after resolution of overt intestinal inflammation. Importantly, we note that circulating exosomes are not exclusively targeted to cardiomyocytes but are capable of delivering functional cargo to multiple organs. Thus, while cardiomyocytes represent one key target in our model, systemic distribution of exosomes may also contribute to broader tissue responses and should be considered when interpreting their role in disease progression.

In summary, our study provides mechanistic evidence that chronic intestinal inflammation drives cardiac dysfunction through a gut microbiota-dependent, GBP1-mediated pathway. These findings identify immune cell mitochondrial remodeling and exosome-mediated signaling as key mechanisms linking colitis to CVD. Beyond DSS induced chronic colitis, this work may also have broader implications for infection-associated cardiac injury, as many pathogens strongly induce GBP family proteins. While GBP1 plays essential antimicrobial roles during infection, excessive or dysregulated GBP1 activation may inadvertently contribute to cardiac dysfunction.

## Nonstandard Abbreviations and Acronyms

ANP: Atrial natriuretic peptide
BMDM: Bone marrow-derived macrophage
BNP: Brain natriuretic peptide
CVD: Cardiovascular disease
DSS: Dextran sulfate sodium
DRP1: Dynamin-related protein 1
ECAR: Extracellular acidification rate
FIS1: Fission 1 protein
FMT: Fecal microbiota transplantation
FS: Fractional shortening
GF: Germ-free
GBP2b/GBP1: Guanylate-binding protein 2b
HSPCs: Hematopoietic stem and progenitor cells
IBD: Inflammatory bowel disease
LCN2: Lipocalin 2
LPS: Lipopolysaccharide
LVEF: Left ventricular ejection fraction
LVIDd: Left ventricular internal diameter at diastole
LVIDs: Left ventricular internal diameter at systole
MYH7: Myosin heavy chain 7
OCR: Oxygen consumption rate
ROS: Reactive oxygen species
TLR4: Toll-like receptor 4
TMAO: Trimethylamine N-oxide

---

**Supplemental Table 1:** Sequences of primers used in the study.

## Supplemental Figure legends

**Figure S1.** Assessment of intestinal inflammation and cardiac alterations in chronic DSS-induced colitis mice. Ten-week-old C57BL/6 mice were treated with 1% DSS in drinking water for one week followed by one week of regular water, repeated over a seven-week period. **A**, Body weight was monitored throughout the experiment. n=5-6 mice/group. **B** & **C**, At the study endpoint, colon length, colon weight-to-length ratio was measured. **D**, Colon tissues were processed for flow cytometric analysis, with the representative gating strategy shown. **E–G,** Frequencies of neutrophils, monocytes, and M1/M2-like macrophage subsets were quantified based on the indicated gating strategy: neutrophils, Ly6G⁺ cells within the CD45⁺MHCII⁻CD11b⁺ population; monocytes, Ly6C^hi/low cells within the CD45⁺MHCII⁻CD11b⁺ population; and M1/M2-like macrophages within the CD45⁺MHCII⁺CD11b⁺F4/80⁺ CD64⁺population. n=5-6 mice/group. **H**, Fecal lipocalin-2 (Lcn2) levels were measured by ELISA. n=5-6 mice/group. **I**, Colon tissues collected at the endpoint were processed for H&E staining. Data are presented as mean ± S.E.M. with statistical significance indicated as follows: * p<0.05, ** p<0.01, *** p<0.001.

**Figure S2.** Chronic DSS-induced colitis causes cardiac remodeling without overt structural damage in major peripheral organs. Ten-week-old C57BL/6 mice were subjected to chronic DSS-induced colitis as described in Figure S1. **A**, Representative images of H&E or picrosirius red (PSR) staining in the mice major organs including kidney, lung, skeletal muscle, epididymal white adipose tissue (eWAT), and liver. **B**, Alanine aminotransferase (ALT) activity in mouse blood was determined using ALT activity assay kit. **C**, Cardiac function in female mice was assessed by echocardiography. n=5 mice/group. **D**, Quantification of ratio of heart weight to tibia length or to mice body weight. n=5 mice/group. **E** through **G**, Total RNA was extracted from heart tissue and subjected to RNA sequencing (RNA-seq). Differential gene expression is shown as a volcano plot (**E**), followed by KEGG pathway enrichment analysis (**F**) and a heatmap of genes associated with mitochondrial ribosomes and ribosome biogenesis (**G**). Data are presented as mean ± S.E.M. with statistical significance indicated as follows: * p<0.05, ** p<0.01, *** p<0.001.

**Figure S3.** DSS-induced chronic colitis does not markedly alter whole-body energy metabolism. **A**, Epididymal fat weight, fat-to-body weight ratio, and serum levels of glucose, cholesterol, triglycerides, and TMAO were assessed in control and DSS induced colitis mice at the end of the experiment. n=5-10 mice/group. **B** through **E**, Control and DSS colitis mice were subjected phenomaster metabolic cage analysis. n=4. **B**, Food and water intake measured by metabolic cage. **C**, Whole body respiratory exchange ratio (RER; VCO2/VO2). **D**, Energy expenditure normalized to body weight. **E**, Locomotor activity. Data are presented as mean ± S.E.M. with statistical significance indicated as follows: * p<0.05, ** p<0.01, *** p<0.001.

**Figure S4.** DSS-induced colitis promotes cardiac dysfunction independent of adaptive immunity. A, Absolute numbers of neutrophils, monocytes, and lymphocytes in the blood of wild-type mice were quantified 7 days after DSS treatment using a hematology cell counter. **B** through **E**, Ten-week-old Rag1 knockout (Rag^-/-^) mice were administered 1% DSS in drinking water for one week, followed by one week of regular water, repeated for total an seven-week period (DSS colitis). n=4-6 mice/group. Control mice received regular drinking water throughout the study (Control). At the end of the experiment, colon length and weight were measured, and the colon weight-to-length ratio was calculated (**B**). Colon tissues were processed for H&E staining to assess histological changes (**C**). Cardiac function was evaluated *via* echocardiography by measuring left ventricular ejection fraction (LVEF) and fractional shortening (FS), and the heart-to-body weight ratio was determined (**D**), and cardiac tissue was analyzed by H&E staining to quantify cardiomyocyte size (**E**). Data are presented as mean ± S.E.M. with statistical significance indicated as follows: * p<0.05, ** p<0.01, *** p<0.001.

**Figure S5.** Persistent immune activation and metabolic reprogramming of circulating immune cells during chronic colitis. **A&B**, circulating innate immune cell populations were analyzed by flow cytometry at day 42 post-DSS. n=5 mice/group. **C** and **D**, OCR and ECAR were measured in circulating immune cells at the later stage of colitis (day 42) using the Seahorse assay. **E**, Heart-infiltrating immune cell populations were analyzed by flow cytometry at day 42 post-DSS. Macrophages were quantified as F4/80⁺ cells within the CD45⁺MHCII⁺CD11b⁺ population, monocytes as Ly6C⁺ cells within the CD45⁺MHCII^-^CD11b⁺Ly6G^-^ population, and neutrophils as Ly6G⁺ cells within the CD45⁺MHCII^-^CD11b⁺ population. **F**, Representative images of H&E staining in the heart of mice, and quantification of percentage of the area of immune cell infiltration. n=5 mice/group. Data are presented as mean ± S.E.M. with statistical significance indicated as follows: * p<0.05, ** p<0.01, *** p<0.001.

**Figure S6.** Persistent systemic reprogramming following colitis and transmission via bone marrow transplantation. **A** through **E**, after a 6-week recovery period following DSS treatment, samples were collected for analysis. **A and B**, colon tissues were collected and processed for H&E staining, and fecal lipocalin-2 levels were measured by ELISA. **C**, The mitochondrial membrane potential of bone marrow stem cells was analyzed by flow cytometry. n=6-8 mice/group. **D** and **E**, Cellular metabolic function was assessed using Seahorse extracellular flux analysis. n=6-8 mice/group. **F** through **J**, Male C57BL/6 mice were lethally irradiated and transplanted with bone marrow cells from either healthy control or DSS colitis donors (**F**). Six weeks after transplantation, the metabolic function of circulating immune cells in recipient mice was evaluated by Seahorse extracellular flux analysis for oxygen consumption rates (OCR), including basal and maximal respiration (**G** and **H**). Cardiac function in recipient mice (**I**) was assessed by echocardiography by measuring ejection fraction (EF) and fractional shortening (FS), and the heart-to-body weight and heart-to-tibia length ratios was determined (**J**). Data are presented as mean ± S.E.M. with statistical significance indicated as follows: n.s., not significant, * p<0.05, ** p<0.01.

**Figure S7.** Dysbiosis persists after colitis and is sufficient to transfer intestinal and systemic phenotypes. **A** and **B**, Fecal samples were collected from control and DSS-treated mice on days 7 and 42 and subjected to 16S rRNA gene sequencing. Overall microbial community structure was analyzed using unweighted UniFrac principal coordinate analysis (PCoA) (**A**). Microbial richness and evenness (α-diversity) were assessed using the Faith’s phylogenetic diversity (PD) index and evenness metrics (**B**). n=6 mice/group. **C**, Fecal samples were collected from control and DSS-treated mice six weeks after DSS withdrawal. Global microbiota composition was evaluated by unweighted UniFrac PCoA, and α-diversity was assessed using the Faith’s PD index and evenness. n=8-10 mice/group. **D–F,** LEfSe (linear discriminant analysis effect size) analysis was performed to identify differentially abundant bacterial genera between DSS-induced colitis mice and control mice at the indicated time points. **G** through I, Germ-free mice received fecal microbiota transplants (FMT) from either healthy control donors or DSS-induced colitis donors. Two months after transplantation, mice were euthanized, and body weight and epididymal fat mass were recorded (**G**). n=5 mice/group. Colonic histopathology was assessed by H&E staining (**H**), and fecal lipocalin-2 (LCN2) levels were measured by ELISA (**I**). Data are presented as mean ± S.E.M. with statistical significance indicated as follows: * p<0.05, ** p<0.01, *** p<0.001.

**Figure S8.** Identification of LPS as the active circulating factor and requirement of hematopoietic TLR4 signaling. **A.** Representative flow cytometry plots showing neutrophils in the heart after anti-Ly6G treatment (ND) or isotype control treatment, and macrophages in the heart after anti-CSF1R plus clodronate liposome treatment (MD) or PBS control treatment. **B.** Quantification of left ventricular ejection fraction (LVEF) and fractional shortening (FS) in mice after neutrophil or macrophage depletion at 28 days post-DSS-induced colitis**. C-E**, Serum from control and DSS-treated mice were subjected to selective biochemical treatments, including heat inactivation, protein digestion (Proteinase K), nucleic acid degradation (DNase/RNase), and lipid extraction. Treated serum were subsequently applied to RAW264.7 macrophages to assess pro-inflammatory activity *via* TNF-α induction. Schematic overview of the workflow used to identify active circulating factors in DSS-induced colitis (**C**). Supernatant TNF-α level after serum treatment (**D**). n=6 mice/group. **E**, RAW264.7 cells were treated with DSS serum in the presence or absence of polymyxin B. Supernatant TNF-α level after serum treatment. n=6 mice/group.

**Figure S9.** Hematopoietic TLR4 signaling mediates microbiota- and LPS-induced immune metabolic reprogramming and cardiac dysfunction. **A**. Serum levels of glucose, cholesterol, and triglycerides, were assessed in Chimeric BM-WT or BM-T4KO mice (generated as shown in Figure 5B) of control and DSS induced colitis mice at the end of the experiment. **B**, Chimeric BM-WT or BM-T4KO mice were colonized with an *E. coli* strain. **C** through **E**, In chimeric mice, circulating immune cell metabolism was evaluated by Seahorse extracellular flux analysis (**C**and **D**), and cardiac function was assessed by echocardiography (**E**). **F** through **H**, Wild-type mice were chronically infused with LPS using osmotic minipumps (**F**). Cardiac function was assessed by echocardiography (**G**), and cardiac weight was analyzed to evaluate hypertrophy (**H**). n=3-5 mice/group. Data are presented as mean ± S.E.M. with statistical significance indicated as follows: n.s., not significant, * p<0.05, ** p<0.01, *** p<0.001.

**Figure S10.** LPS-TLR4 signaling induces GBP1 expression and mitochondrial localization in immune cells. **A**, Quantitative RT-PCR analysis of guanylate-binding proteins (GBPs) expression in circulating immune cells at day 42 post-DSS treatment. n=6 mice/group. **B**, GBP1 expression in heart tissue was assessed by immunohistochemistry, and GBP1-positive area was quantified at day 42 post-DSS treatment. n=6 mice/group. **C**, Heart sections were stained for CD68, GBP1, and DAPI, and the colocalization of CD68 with GBP1 was quantified. n=6 mice/group. **D**, Subcellular fractionation of THP-1 monocytes following LPS stimulation, followed by immunoblot analysis of GBP1 in whole-cell, cytosolic, and mitochondrial fractions. **E**, Confocal microscopy of THP-1 cells stained for GBP1 and the mitochondrial marker FIS1 to assess their colocalization after LPS treatment. n=9. Data are presented as mean ± S.E.M. with statistical significance indicated as follows: n.s., not significant, ** p<0.01, *** p<0.001.

**Figure S11.** Systemic biodistribution and cardiomyocyte uptake of circulating exosomes. **A** through **C**, DiR-labeled exosomes (Exo-DiR) were administered to mice *via* tail-vein injection. Control mice received an equivalent amount of PBS or free DiR dye (Free-DiR) were processed under identical conditions. *In vivo* fluorescence signals of mice whole body after 2- or 24-h injection (**A**) and *ex vivo* fluorescence images of major organs from mice 24 hours post injection (**B**) were detected by IVIS system, and fluorescence signals (red) in the heart were further confirmed by staining with DAPI (blue) in the frozen sections (**C**). **D**, H9C2 cardiomyocytes were treated with control or DiR-labeled exosomes for 24 h, DiR signals (red) and Hoechst-stained nucleus (blue) were captured by a fluorescent microscopy.

**Figure S12.** GBP1-containing exosomes in colitis mediate cardiomyocyte hypertrophy and cardiac fibrosis. **A**, Wild-type mice were intravenously injected with exosomes isolated from serum of control or DSS-induced colitis mice. Cardiac fibrosis was assessed by Picrosirius Red (PSR) staining, and collagen content was quantified. n=5 mice/group. **B**, THP-1 cells were treated with LPS with or without GBP1 silencing. GBP1 expression in cells and secreted exosomes was analyzed by immunoblotting. **C** through **E**, Exosomes isolated from vehicle- or LPS-treated THP-1 cells, with or without GBP1 silencing, were used to treat H9c2 cardiomyocytes. Cardiomyocyte hypertrophy was assessed by measuring cell surface area, with representative images shown (**C**) and quantitative analyses of total protein content (**D**) and cell size (**E**). n= 6. **F**, Wild-type mice were intravenously injected with exosomes isolated from THP-1 cells treated with vehicle or LPS, with or without GBP1 silencing. Cardiac fibrosis was evaluated by PSR staining, and collagen content was quantified. n=5-6 mice/group. Data are presented as mean ± S.E.M. with statistical significance indicated as follows: * p<0.05, ** p<0.01, *** p<0.001.

## REFERENCES

1. Coronado F, Melvin SC, Bell RA, Zhao G. Global Responses to Prevent, Manage, and Control Cardiovascular Diseases. Prev Chronic Dis. 2022;19:E84. doi: 10.5888/pcd19.220347

2. Roth GA, Mensah GA, Johnson CO, Addolorato G, Ammirati E, Baddour LM, Barengo NC, Beaton AZ, Benjamin EJ, Benziger CP, et al. Global Burden of Cardiovascular Diseases and Risk Factors, 1990-2019: Update From the GBD 2019 Study. J Am Coll Cardiol. 2020;76:2982–3021. doi: 10.1016/j.jacc.2020.11.010

3. Sun J, Qiao Y, Zhao M, Magnussen CG, Xi B. Global, regional, and national burden of cardiovascular diseases in youths and young adults aged 15-39 years in 204 countries/territories, 1990-2019: a systematic analysis of Global Burden of Disease Study 2019. BMC Med. 2023;21:222. doi: 10.1186/s12916-023-02925-4

4. Wu H, Hu T, Hao H, Hill MA, Xu C, Liu Z. Inflammatory bowel disease and cardiovascular diseases: a concise review. Eur Heart J Open. 2022;2:oeab029. doi: 10.1093/ehjopen/oeab029

5. Aniwan S, Pardi DS, Tremaine WJ, Loftus EV, Jr. Increased Risk of Acute Myocardial Infarction and Heart Failure in Patients With Inflammatory Bowel Diseases. Clin Gastroenterol Hepatol. 2018;16:1607–1615 e1601. doi: 10.1016/j.cgh.2018.04.031

6. Czubkowski P, Osiecki M, Szymanska E, Kierkus J. The risk of cardiovascular complications in inflammatory bowel disease. Clin Exp Med. 2020;20:481–491. doi: 10.1007/s10238-020-00639-y

7. Bernstein CN, Wajda A, Blanchard JF. The incidence of arterial thromboembolic diseases in inflammatory bowel disease: a population-based study. Clin Gastroenterol Hepatol. 2008;6:41–45. doi: 10.1016/j.cgh.2007.09.016

8. Rungoe C, Basit S, Ranthe MF, Wohlfahrt J, Langholz E, Jess T. Risk of ischaemic heart disease in patients with inflammatory bowel disease: a nationwide Danish cohort study. Gut. 2013;62:689–694. doi: 10.1136/gutjnl-2012-303285

9. Xavier RJ, Podolsky DK. Unravelling the pathogenesis of inflammatory bowel disease. Nature. 2007;448:427–434. doi: 10.1038/nature06005

10. Turner JR. Intestinal mucosal barrier function in health and disease. Nat Rev Immunol. 2009;9:799–809. doi: 10.1038/nri2653

11. Rogler G, Singh A, Kavanaugh A, Rubin DT. Extraintestinal Manifestations of Inflammatory Bowel Disease: Current Concepts, Treatment, and Implications for Disease Management. Gastroenterology. 2021;161:1118–1132. doi: 10.1053/j.gastro.2021.07.042

12. Witkowski M, Weeks TL, Hazen SL. Gut Microbiota and Cardiovascular Disease. Circ Res. 2020;127:553–570. doi: 10.1161/CIRCRESAHA.120.316242

13. Li J, Wang Y, Shrestha S, Gewirtz AT, Ding Y, Zou J. Targeting Gut Microbiota to Combat Vascular Aging and Cardiovascular Disease: Mechanisms and Therapeutic Potential. Nutrients. 2025;17. doi: 10.3390/nu17172887

14. Kostic AD, Xavier RJ, Gevers D. The microbiome in inflammatory bowel disease: current status and the future ahead. Gastroenterology. 2014;146:1489–1499. doi: 10.1053/j.gastro.2014.02.009

15. Gevers D, Kugathasan S, Denson LA, Vazquez-Baeza Y, Van Treuren W, Ren B, Schwager E, Knights D, Song SJ, Yassour M, et al. The treatment-naive microbiome in new-onset Crohn’s disease. Cell Host Microbe. 2014;15:382–392. doi: 10.1016/j.chom.2014.02.005

16. Zhao X, Wang Y, Wang Y, Gewirtz AT, Zou J. Inulin aggravates colitis through gut microbiota modulation and MyD88/IL-18 signaling. Gut Microbes. 2025;17:2570425. doi: 10.1080/19490976.2025.2570425

17. Kirkby M, Enosi Tuipulotu D, Feng S, Lo Pilato J, Man SM. Guanylate-binding proteins: mechanisms of pattern recognition and antimicrobial functions. Trends Biochem Sci. 2023;48:883–893. doi: 10.1016/j.tibs.2023.07.002

18. Feng S, Enosi Tuipulotu D, Pandey A, Jing W, Shen C, Ngo C, Tessema MB, Li FJ, Fox D, Mathur A, et al. Pathogen-selective killing by guanylate-binding proteins as a molecular mechanism leading to inflammasome signaling. Nat Commun. 2022;13:4395. doi: 10.1038/s41467-022-32127-0

19. Tailor D, Garcia-Marques FJ, Bermudez A, Pitteri SJ, Malhotra SV. Guanylate-binding protein 1 modulates proteasomal machinery in ovarian cancer. iScience. 2023;26:108292. doi: 10.1016/j.isci.2023.108292

20. Honkala AT, Tailor D, Malhotra SV. Guanylate-Binding Protein 1: An Emerging Target in Inflammation and Cancer. Front Immunol. 2019;10:3139. doi: 10.3389/fimmu.2019.03139

21. Xie J-H, Li Y-Y, Jin J. The essential functions of mitochondrial dynamics in immune cells. Cellular & molecular immunology. 2020;17:712–721.

22. Rambold AS, Pearce EL. Mitochondrial dynamics at the interface of immune cell metabolism and function. Trends in immunology. 2018;39:6–18.

23. Bao D, Zhao J, Zhou X, Yang Q, Chen Y, Zhu J, Yuan P, Yang J, Qin T, Wan S. Mitochondrial fission-induced mtDNA stress promotes tumor-associated macrophage infiltration and HCC progression. Oncogene. 2019;38:5007–5020.

24. Marques E, Kramer R, Ryan DG. Multifaceted mitochondria in innate immunity. npj Metabolic Health and Disease. 2024;2:6.

25. Mohanty A, Tiwari-Pandey R, Pandey NR. Mitochondria: the indispensable players in innate immunity and guardians of the inflammatory response. Journal of cell communication and signaling. 2019;13:303–318.

26. Angajala A, Lim S, Phillips JB, Kim J-H, Yates C, You Z, Tan M. Diverse roles of mitochondria in immune responses: novel insights into immuno-metabolism. Frontiers in immunology. 2018;9:1605.

27. West AP, Shadel GS, Ghosh S. Mitochondria in innate immune responses. Nature Reviews Immunology. 2011;11:389–402.

28. Forte M, Schirone L, Ameri P, Basso C, Catalucci D, Modica J, Chimenti C, Crotti L, Frati G, Rubattu S. The role of mitochondrial dynamics in cardiovascular diseases. British journal of pharmacology. 2021;178:2060–2076.

29. Cruz CSD, Kang M-J. Mitochondrial dysfunction and damage associated molecular patterns (DAMPs) in chronic inflammatory diseases. Mitochondrion. 2018;41:37–44.

30. Li J, Li D, Zhao F, Wang Y, Yao H, Wu Y, An J, Liu Z, Ding Y, Zou MH. Endothelial FUNDC1 regulates metabolic reprogramming and the obesity-diabetes transition through the SIRT3/GATA2/endothelin-1 axis. Nat Commun. 2026;17:1836. doi: 10.1038/s41467-026-68548-4

31. Zou J, Ngo VL, Wang Y, Wang Y, Gewirtz AT. Maternal fiber deprivation alters microbiota in offspring, resulting in low-grade inflammation and predisposition to obesity. Cell Host Microbe. 2023;31:45–57 e47. doi: 10.1016/j.chom.2022.10.014

32. Monteiro LB, Davanzo GG, de Aguiar CF, Moraes-Vieira PMM. Using flow cytometry for mitochondrial assays. MethodsX. 2020;7:100938. doi: 10.1016/j.mex.2020.100938

33. Cani PD, Amar J, Iglesias MA, Poggi M, Knauf C, Bastelica D, Neyrinck AM, Fava F, Tuohy KM, Chabo C, et al. Metabolic endotoxemia initiates obesity and insulin resistance. Diabetes. 2007;56:1761–1772. doi: 10.2337/db06-1491

34. Sun J, Yao J, Olen O, Halfvarson J, Bergman D, Ebrahimi F, Rosengren A, Sundstrom J, Ludvigsson JF. Risk of heart failure in inflammatory bowel disease: a Swedish population-based study. Eur Heart J. 2024;45:2493–2504. doi: 10.1093/eurheartj/ehae338

35. Zakerska-Banaszak O, Tomczak H, Gabryel M, Baturo A, Wolko L, Michalak M, Malinska N, Mankowska-Wierzbicka D, Eder P, Dobrowolska A, et al. Dysbiosis of gut microbiota in Polish patients with ulcerative colitis: a pilot study. Sci Rep. 2021;11:2166. doi: 10.1038/s41598-021-81628-3

36. Carrillo-Salinas FJ, Anastasiou M, Ngwenyama N, Kaur K, Tai A, Smolgovsky SA, Jetton D, Aronovitz M, Alcaide P. Gut dysbiosis induced by cardiac pressure overload enhances adverse cardiac remodeling in a T cell-dependent manner. Gut Microbes. 2020;12:1–20. doi: 10.1080/19490976.2020.1823801

37. Mirsepasi-Lauridsen HC, Vallance BA, Krogfelt KA, Petersen AM. Escherichia coli Pathobionts Associated with Inflammatory Bowel Disease. Clin Microbiol Rev. 2019;32. doi: 10.1128/CMR.00060-18

38. Yu T, Wang L, Zhang L, Deuster PA. Mitochondrial Fission as a Therapeutic Target for Metabolic Diseases: Insights into Antioxidant Strategies. Antioxidants (Basel). 2023;12. doi: 10.3390/antiox12061163

39. Chen M, Wu Y, Chen C. Extracellular Vesicles as Emerging Regulators in Ischemic and Hypertrophic Cardiovascular Diseases: A Review of Pathogenesis and Therapeutics. Med Sci Monit. 2025;31:e948948. doi: 10.12659/MSM.948948

40. Fumery M, Singh S, Dulai PS, Gower-Rousseau C, Peyrin-Biroulet L, Sandborn WJ. Natural History of Adult Ulcerative Colitis in Population-based Cohorts: A Systematic Review. Clin Gastroenterol Hepatol. 2018;16:343–356 e343. doi: 10.1016/j.cgh.2017.06.016

41. Weimers P, Munkholm P. The Natural History of IBD: Lessons Learned. Curr Treat Options Gastroenterol. 2018;16:101–111. doi: 10.1007/s11938-018-0173-3

42. Baars JE, Nuij VJ, Oldenburg B, Kuipers EJ, van der Woude CJ. Majority of patients with inflammatory bowel disease in clinical remission have mucosal inflammation. Inflamm Bowel Dis. 2012;18:1634–1640. doi: 10.1002/ibd.21925

43. Park S, Abdi T, Gentry M, Laine L. Histological Disease Activity as a Predictor of Clinical Relapse Among Patients With Ulcerative Colitis: Systematic Review and Meta-Analysis. Am J Gastroenterol. 2016;111:1692–1701. doi: 10.1038/ajg.2016.418

44. Khorsand B, Asadzadeh Aghdaei H, Nazemalhosseini-Mojarad E, Nadalian B, Nadalian B, Houri H. Overrepresentation of Enterobacteriaceae and Escherichia coli is the major gut microbiome signature in Crohn’s disease and ulcerative colitis; a comprehensive metagenomic analysis of IBDMDB datasets. Front Cell Infect Microbiol. 2022;12:1015890. doi: 10.3389/fcimb.2022.1015890

45. Fang X, Monk JM, Nurk S, Akseshina M, Zhu Q, Gemmell C, Gianetto-Hill C, Leung N, Szubin R, Sanders J, et al. Metagenomics-Based, Strain-Level Analysis of Escherichia coli From a Time-Series of Microbiome Samples From a Crohn’s Disease Patient. Front Microbiol. 2018;9:2559. doi: 10.3389/fmicb.2018.02559

46. Kondapalli N, Katari V, Dalal KK, Paruchuri S, Thodeti CK. Microbiota in Gut-Heart Axis: Metabolites and Mechanisms in Cardiovascular Disease. Compr Physiol. 2025;15:e70024. doi: 10.1002/cph4.70024

47. Berger M, Kleber ME, Delgado GE, Marz W, Andreas M, Hellstern P, Marx N, Schuett KA. Trimethylamine N-Oxide and Adenosine Diphosphate-Induced Platelet Reactivity Are Independent Risk Factors for Cardiovascular and All-Cause Mortality. Circ Res. 2020;126:660–662. doi: 10.1161/CIRCRESAHA.119.316214

48. Lepper PM, Kleber ME, Grammer TB, Hoffmann K, Dietz S, Winkelmann BR, Boehm BO, Marz W. Lipopolysaccharide-binding protein (LBP) is associated with total and cardiovascular mortality in individuals with or without stable coronary artery disease--results from the Ludwigshafen Risk and Cardiovascular Health Study (LURIC). Atherosclerosis. 2011;219:291–297. doi: 10.1016/j.atherosclerosis.2011.06.001

49. Singh MV, Cicha MZ, Nunez S, Meyerholz DK, Chapleau MW, Abboud FM. Angiotensin II-induced hypertension and cardiac hypertrophy are differentially mediated by TLR3- and TLR4-dependent pathways. Am J Physiol Heart Circ Physiol. 2019;316:H1027–H1038. doi: 10.1152/ajpheart.00697.2018

50. Singh MV, Cicha MZ, Meyerholz DK, Chapleau MW, Abboud FM. Dual Activation of TRIF and MyD88 Adaptor Proteins by Angiotensin II Evokes Opposing Effects on Pressure, Cardiac Hypertrophy, and Inflammatory Gene Expression. Hypertension. 2015;66:647–656. doi: 10.1161/HYPERTENSIONAHA.115.06011

51. Mian MOR, He Y, Bertagnolli M, Mai-Vo TA, Fernandes RO, Boudreau F, Cloutier A, Luu TM, Nuyt AM. TLR (Toll-Like Receptor) 4 Antagonism Prevents Left Ventricular Hypertrophy and Dysfunction Caused by Neonatal Hyperoxia Exposure in Rats. Hypertension. 2019;74:843–853. doi: 10.1161/HYPERTENSIONAHA.119.13022

52. Wiger CW, Ranheim T, Arnesen H, Vaage J, Pischke SE, Yndestad A, Stenslokken KO, Torp MK. TLR4 Inhibition Attenuated LPS-Induced Proinflammatory Signaling and Cytokine Release in Mouse Hearts and Cardiomyocytes. Immun Inflamm Dis. 2025;13:e70133. doi: 10.1002/iid3.70133

53. Panaro MA, Gagliardi N, Saponaro C, Calvello R, Mitolo V, Cianciulli A. Toll-like receptor 4 mediates LPS-induced release of nitric oxide and tumor necrosis factor-alpha by embryonal cardiomyocytes: biological significance and clinical implications in human pathology. Curr Pharm Des. 2010;16:766–774. doi: 10.2174/138161210790883624

54. Gabele E, Dostert K, Hofmann C, Wiest R, Scholmerich J, Hellerbrand C, Obermeier F. DSS induced colitis increases portal LPS levels and enhances hepatic inflammation and fibrogenesis in experimental NASH. J Hepatol. 2011;55:1391–1399. doi: 10.1016/j.jhep.2011.02.035

55. Chait A, den Hartigh LJ. Adipose Tissue Distribution, Inflammation and Its Metabolic Consequences, Including Diabetes and Cardiovascular Disease. Front Cardiovasc Med. 2020;7:22. doi: 10.3389/fcvm.2020.00022

56. Theofilis P, Mystakidi VC, Goliopoulou A, Papamikroulis GA, Lazaros G, Anastasiou M, Tsalamandris S, Vavouranaki G, Korakas E, Lambadiari V, et al. Hepatic steatosis and its association with left ventricular concentric remodeling: insights from the Corinthia study. Hellenic J Cardiol. 2025;85:108–110. doi: 10.1016/j.hjc.2024.10.007

57. Marchetti P, Fovez Q, Germain N, Khamari R, Kluza J. Mitochondrial spare respiratory capacity: mechanisms, regulation, and significance in non-transformed and cancer cells. The FASEB Journal. 2020;34:13106–13124.

58. Flynn JM, Choi SW, Day NU, Gerencser AA, Hubbard A, Melov S. Impaired spare respiratory capacity in cortical synaptosomes from Sod2 null mice. Free Radical Biology and Medicine. 2011;50:866–873.

59. Katkenov N, Mukhatayev Z, Kozhakhmetov S, Sailybayeva A, Bekbossynova M, Kushugulova A. Systematic Review on the Role of IL-6 and IL-1beta in Cardiovascular Diseases. J Cardiovasc Dev Dis. 2024;11. doi: 10.3390/jcdd11070206

60. Feng Y, Ye D, Wang Z, Pan H, Lu X, Wang M, Xu Y, Yu J, Zhang J, Zhao M, et al. The Role of Interleukin-6 Family Members in Cardiovascular Diseases. Front Cardiovasc Med. 2022;9:818890. doi: 10.3389/fcvm.2022.818890

61. Kanda T, Takahashi T. Interleukin-6 and cardiovascular diseases. Jpn Heart J. 2004;45:183–193. doi: 10.1536/jhj.45.183

62. Finethy R, Jorgensen I, Haldar AK, de Zoete MR, Strowig T, Flavell RA, Yamamoto M, Nagarajan UM, Miao EA, Coers J. Guanylate binding proteins enable rapid activation of canonical and noncanonical inflammasomes in Chlamydia-infected macrophages. Infect Immun. 2015;83:4740–4749. doi: 10.1128/IAI.00856-15

63. Waldenstrom A, Ronquist G. Role of exosomes in myocardial remodeling. Circ Res. 2014;114:315–324. doi: 10.1161/CIRCRESAHA.114.300584

64. Ranjan P, Colin K, Dutta RK, Verma SK. Challenges and future scope of exosomes in the treatment of cardiovascular diseases. J Physiol. 2023;601:4873–4893. doi: 10.1113/JP282053

65. Liu C, Wu R, Yang H, Yao Y. Immune cell dynamics and their role in cardiac injury: Mechanisms and therapeutic implications. Biomedicine & Pharmacotherapy. 2025;192:118608.

66. Wen H, Peng L, Chen Y. The effect of immune cell-derived exosomes in the cardiac tissue repair after myocardial infarction: Molecular mechanisms and pre-clinical evidence. Journal of Cellular and Molecular Medicine. 2021;25:6500–6510.

