## Supplemental Figures for "Gut Microbiota Dysbiosis Drives Myocardial Hypertrophy Through GBP2b/GBP1-Mediated Immune Reprogramming and Exosomal Signaling in Chronic Colitis"

**Supplemental Table 1. Sequences of primers used in the study**

| Species | Gene name | Primer Sequence (Forward) | Primer Sequence (Reverse) |
|---|---|---|---|
| **Mouse** | ***Nppa*** | GCTTCCAGGCCATATTGGAG | GGGGGCATGACCTCATCTT |
| | ***Nppb*** | GAGGTCACTCCTATCCTCTGG | GCCATTTCCTCCGACTTTTCTC |
| | ***Myh7*** | ACTGTCAACACTAAGAGGGTCA | TTGGATGATTTGATCTTCCAGGG |
| | ***Gapdh*** | AGGTCGGTGTGAACGGATTTG | TGTAGACCATGTAGTTGAGGTCA |
| | ***Gbp1*** | ACAACTCAGCTAACTTTGTGGG | TGATACACAGGCGAGGCATATTA |
| | ***Gbp2*** | CTGCACTATGTGACGGAGCTA | GAGTCCACACAAAGGTTGGAAA |
| | ***Gbp3*** | GAGGCACCCATTTGTCTGGT | CCGTCCTGCAAGACGATTCA |
| | ***Gbp4*** | GGAGAAGCTAACGAAGGAACAA | TTCCACAAGGGAATCACCATTTT |
| | ***Gbp5*** | CAGACCTATTTGAACGCCAAAGA | TGCCTTGATTCTATCAGCCTCT |
| | ***Gbp6*** | GTTCCAGGAAGTAACAAAGGCT | ATCCCTAGTCTATTCCCAGTGAC |
| | ***Gbp7*** | TCCTGTGTGCCTAGTGGAAAA | CAAGCGGTTCATCAAGTAGGAT |
| **Rat** | ***Nppa*** | GGAGCCTGCGAAGGTCAA | TATCTTCGGTACCGGAAGCTGT |
| | ***Nppb*** | CAGAAGCTGGAGCTGATAAG | TGTAGGGCCTTGGTCCTTTG |
| | ***Myh7*** | GCTGTTATTGCAGCCATTG | TTCCTGTTGCCCCAAAATG |
| | ***Gapdh*** | ACAGCAACAGGGTGGTGGAC | TTTGAGGGTGCAGCGAACTT |

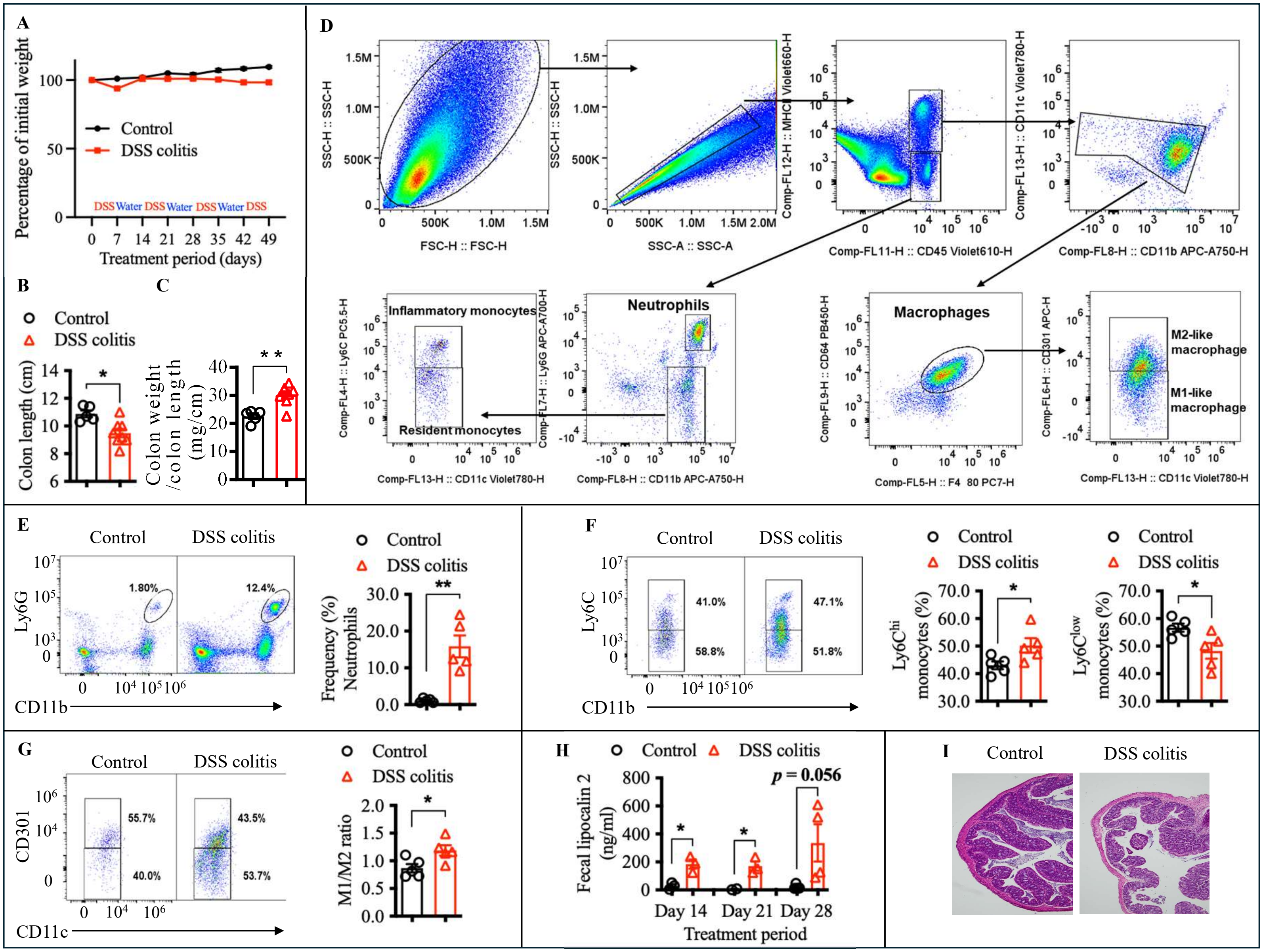


**Figure S1. Assessment of intestinal inflammation and cardiac alterations in chronic DSS-induced colitis mice.** Ten-week-old C57BL/6 mice were treated with 1% DSS in drinking water for one week followed by one week of regular water, repeated over a seven-week period. **A**, Body weight was monitored throughout the experiment. n=5-6 mice/group. **B** & **C**, At the study endpoint, colon length, colon weight-to-length ratio was measured. D, Colon tissues were processed for flow cytometric analysis, with the representative gating strategy shown. E–G, Frequencies of neutrophils, monocytes, and M1/M2-like macrophage subsets were quantified based on the indicated gating strategy: neutrophils, $Ly6G^+$ cells within the $CD45^+MHCII^-CD11b^+$ population; monocytes, Ly6C^hi/low cells within the $CD45^+MHCII^-CD11b^+$ population; and M1/M2-like macrophages within the $CD45^+MHCII^+CD11b^+F4/80^+$ $CD64^+$population. n=5-6 mice/group. **H**, Fecal lipocalin-2 (Lcn2) levels were measured by ELISA. n=5-6 mice/group. **I**, Colon tissues collected at the endpoint were processed for H&E staining. Data are presented as mean ± S.E.M. with statistical significance indicated as follows: * p<0.05, ** p<0.01, *** p<0.001.

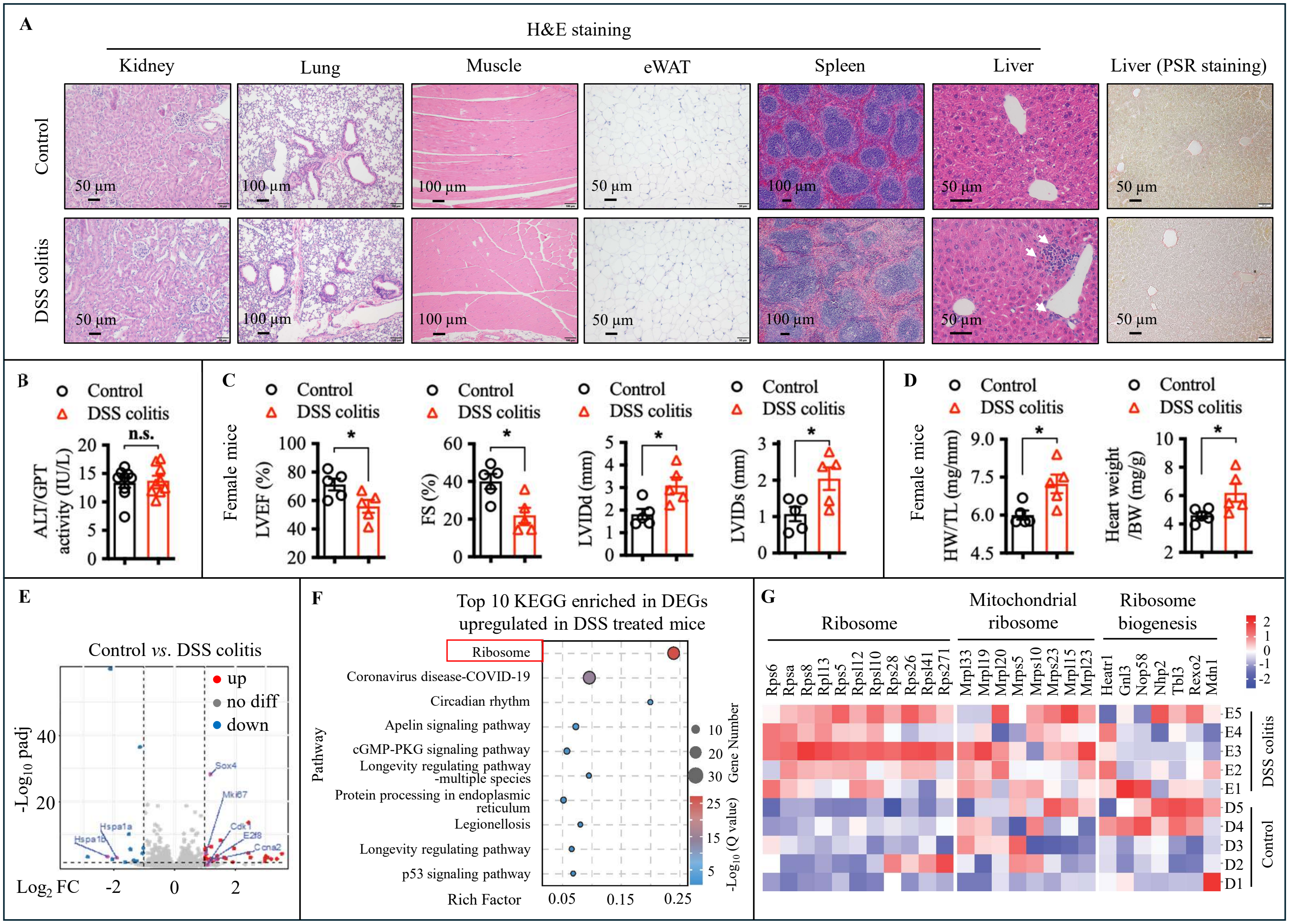


**Figure S2. Chronic DSS-induced colitis causes cardiac remodeling without overt structural damage in major peripheral organs.** Ten-week-old C57BL/6 mice were subjected to chronic DSS-induced colitis as described in Figure S1. **A**, Representative images of H&E or picrosirius red (PSR) staining in the mice major organs including kidney, lung, skeletal muscle, epididymal white adipose tissue (eWAT), and liver. **B**, Alanine aminotransferase (ALT) activity in mouse blood was determined using ALT activity assay kit. **C**, Cardiac function in female mice was assessed by echocardiography. n=5 mice/group. **D**, Quantification of ratio of heart weight to tibia length or to mice body weight. n=5 mice/group. **E** through **G**, Total RNA was extracted from heart tissue and subjected to RNA sequencing (RNA-seq). Differential gene expression is shown as a volcano plot (**E**), followed by KEGG pathway enrichment analysis (**F**) and a heatmap of genes associated with mitochondrial ribosomes and ribosome biogenesis (**G**). Data are presented as mean ± S.E.M. with statistical significance indicated as follows: * p<0.05, ** p<0.01, *** p<0.001.

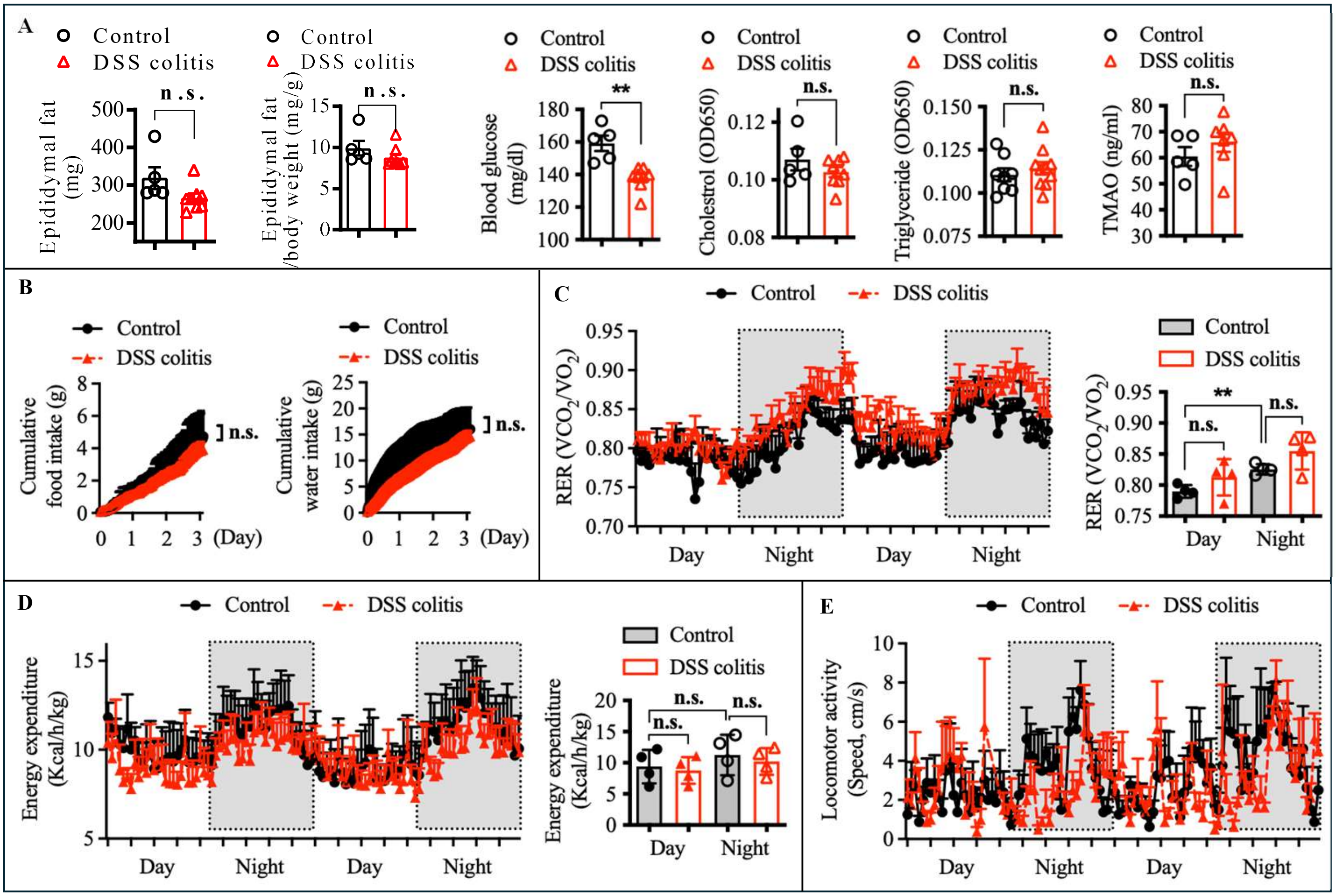


**Figure S3. DSS-induced chronic colitis does not markedly alter whole-body energy metabolism. A**, Epididymal fat weight, fat-to-body weight ratio, and serum levels of glucose, cholesterol, triglycerides, and TMAO were assessed in control and DSS induced colitis mice at the end of the experiment. n=5-10 mice/group. **B** through **E**, Control and DSS colitis mice were subjected phenomaster metabolic cage analysis. n=4. **B**, Food and water intake measured by metabolic cage. **C**, Whole body respiratory exchange ratio (RER; $VCO_2/VO_2$). **D**, Energy expenditure normalized to body weight. **E**, Locomotor activity. Data are presented as mean ± S.E.M. with statistical significance indicated as follows: * p<0.05, ** p<0.01, *** p<0.001.

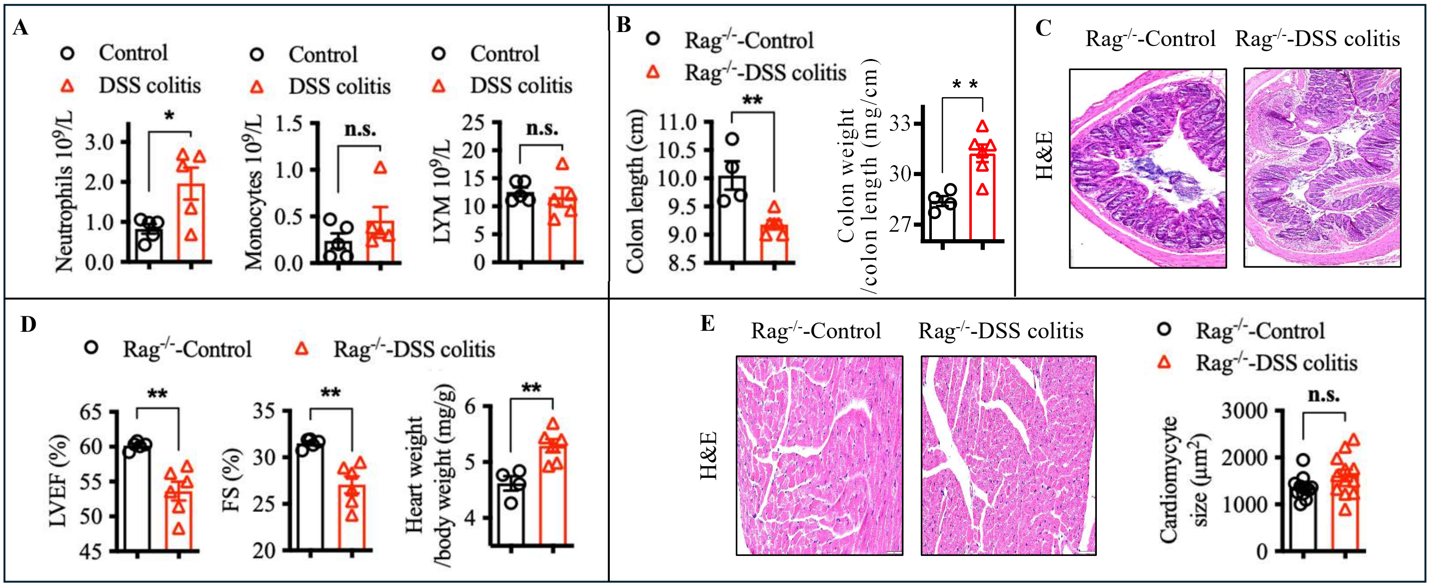


**Figure S4. DSS-induced colitis promotes cardiac dysfunction independent of adaptive immunity. A,** Absolute numbers of neutrophils, monocytes, and lymphocytes in the blood of wild-type mice were quantified 7 days after DSS treatment using a hematology cell counter. **B** through **E**, Ten-week-old Rag1 knockout ($Rag^{-/-}$) mice were administered 1% DSS in drinking water for one week, followed by one week of regular water, repeated for total an seven-week period (DSS colitis). n=4-6 mice/group. Control mice received regular drinking water throughout the study (Control). At the end of the experiment, colon length and weight were measured, and the colon weight-to-length ratio was calculated (**B**). Colon tissues were processed for H&E staining to assess histological changes (**C**). Cardiac function was evaluated *via* echocardiography by measuring left ventricular ejection fraction (LVEF) and fractional shortening (FS), and the heart-to-body weight ratio was determined (**D**), and cardiac tissue was analyzed by H&E staining to quantify cardiomyocyte size (**E**). Data are presented as mean ± S.E.M. with statistical significance indicated as follows: * $p<0.05$, ** $p<0.01$, *** $p<0.001$.

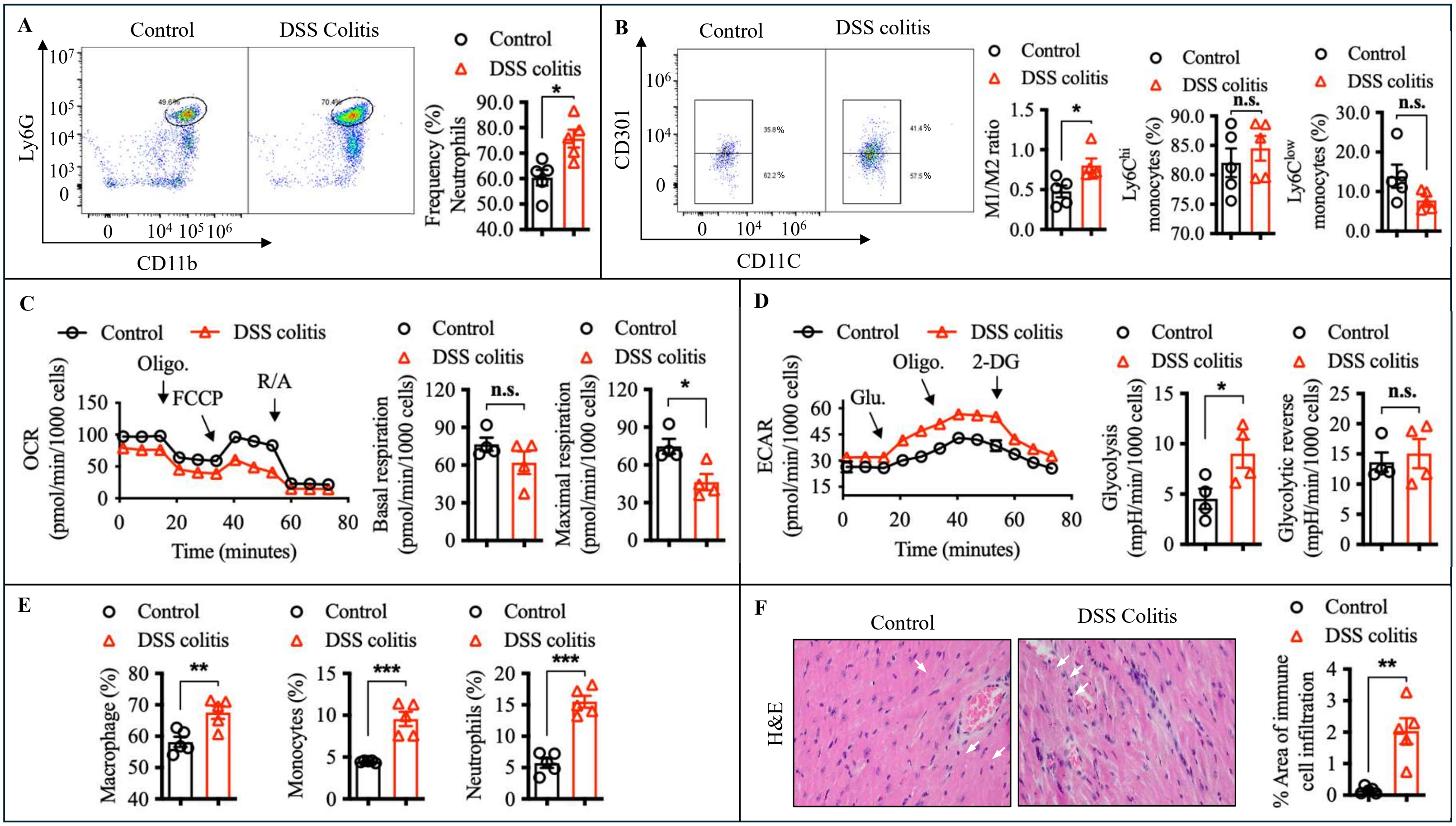

**Figure S5. Persistent immune activation and metabolic reprogramming of circulating immune cells during chronic colitis. A&B**, circulating innate immune cell populations were analyzed by flow cytometry at day 42 post-DSS. n=5 mice/group. **C** and **D**, OCR and ECAR were measured in circulating immune cells at the later stage of colitis (day 42) using the Seahorse assay. **E**, Heart-infiltrating immune cell populations were analyzed by flow cytometry at day 42 post-DSS. Macrophages were quantified as F4/80$^+$ cells within the CD45$^+$MHCII$^+$CD11b$^+$ population, monocytes as Ly6C$^+$ cells within the CD45$^+$MHCII$^-$CD11b$^+$Ly6G$^-$ population, and neutrophils as Ly6G$^+$ cells within the CD45$^+$MHCII$^-$CD11b$^+$ population. **F**, Representative images of H&E staining in the heart of mice, and quantification of percentage of the area of immune cell infiltration. n=5 mice/group. Data are presented as mean ± S.E.M. with statistical significance indicated as follows: * p<0.05, ** p<0.01, *** p<0.001.

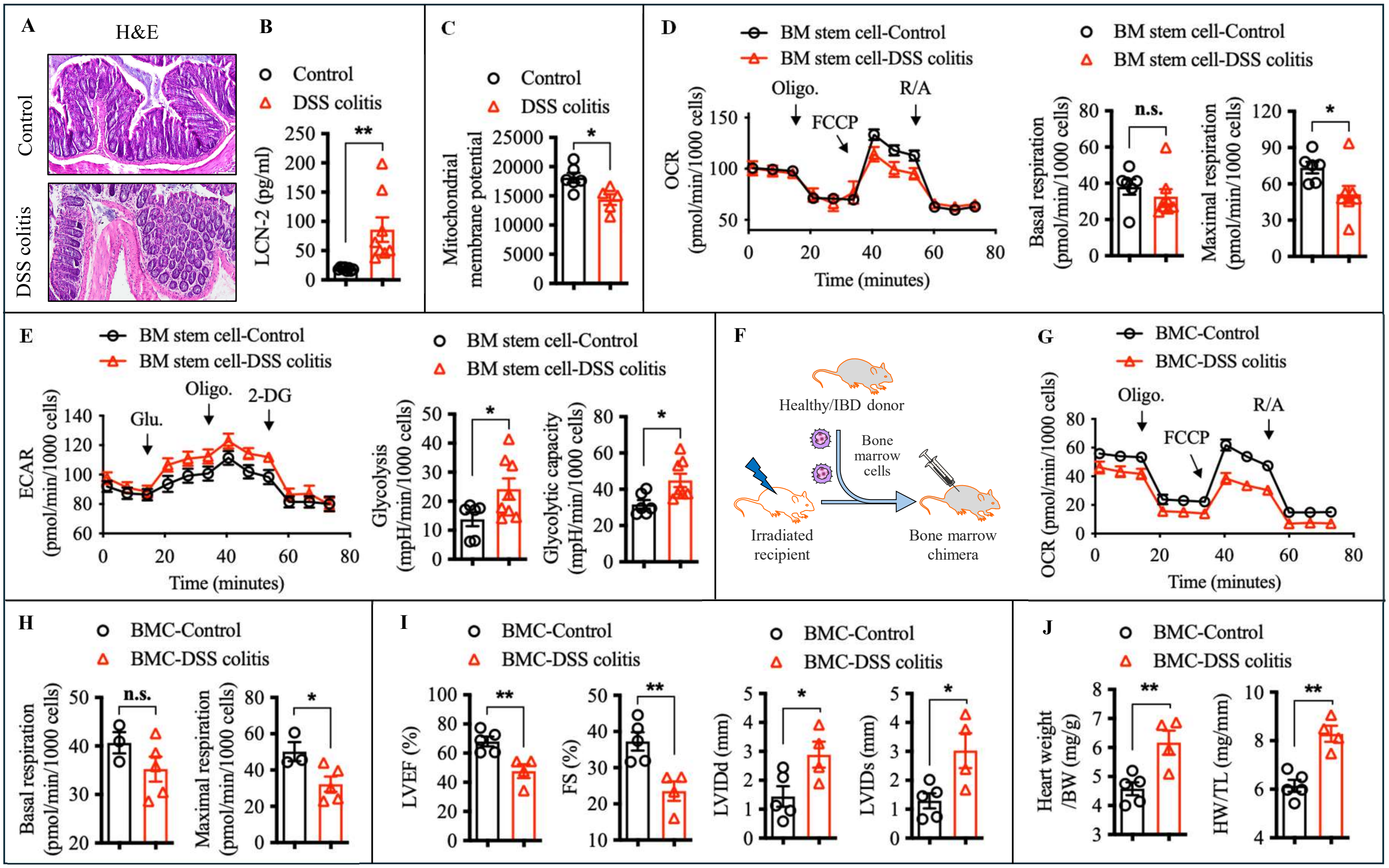


**Figure S6. Persistent systemic reprogramming following colitis and transmission via bone marrow transplantation.** **A** through **E**, after a 6-week recovery period following DSS treatment, samples were collected for analysis. **A and B**, colon tissues were collected and processed for H&E staining, and fecal lipocalin-2 levels were measured by ELISA. **C**, The mitochondrial membrane potential of bone marrow stem cells was analyzed by flow cytometry. n=6-8 mice/group. **D** and **E**, Cellular metabolic function was assessed using Seahorse extracellular flux analysis. n=6-8 mice/group. **F** through **J**, Male C57BL/6 mice were lethally irradiated and transplanted with bone marrow cells from either healthy control or DSS colitis donors (**F**). Six weeks after transplantation, the metabolic function of circulating immune cells in recipient mice was evaluated by Seahorse extracellular flux analysis for oxygen consumption rates (OCR), including basal and maximal respiration (**G** and **H**). Cardiac function in recipient mice (**I**) was assessed by echocardiography by measuring ejection fraction (EF) and fractional shortening (FS), and the heart-to-body weight and heart-to-tibia length ratios was determined (**J**). Data are presented as mean ± S.E.M. with statistical significance indicated as follows: n.s., not significant, * $p<0.05$, ** $p<0.01$.

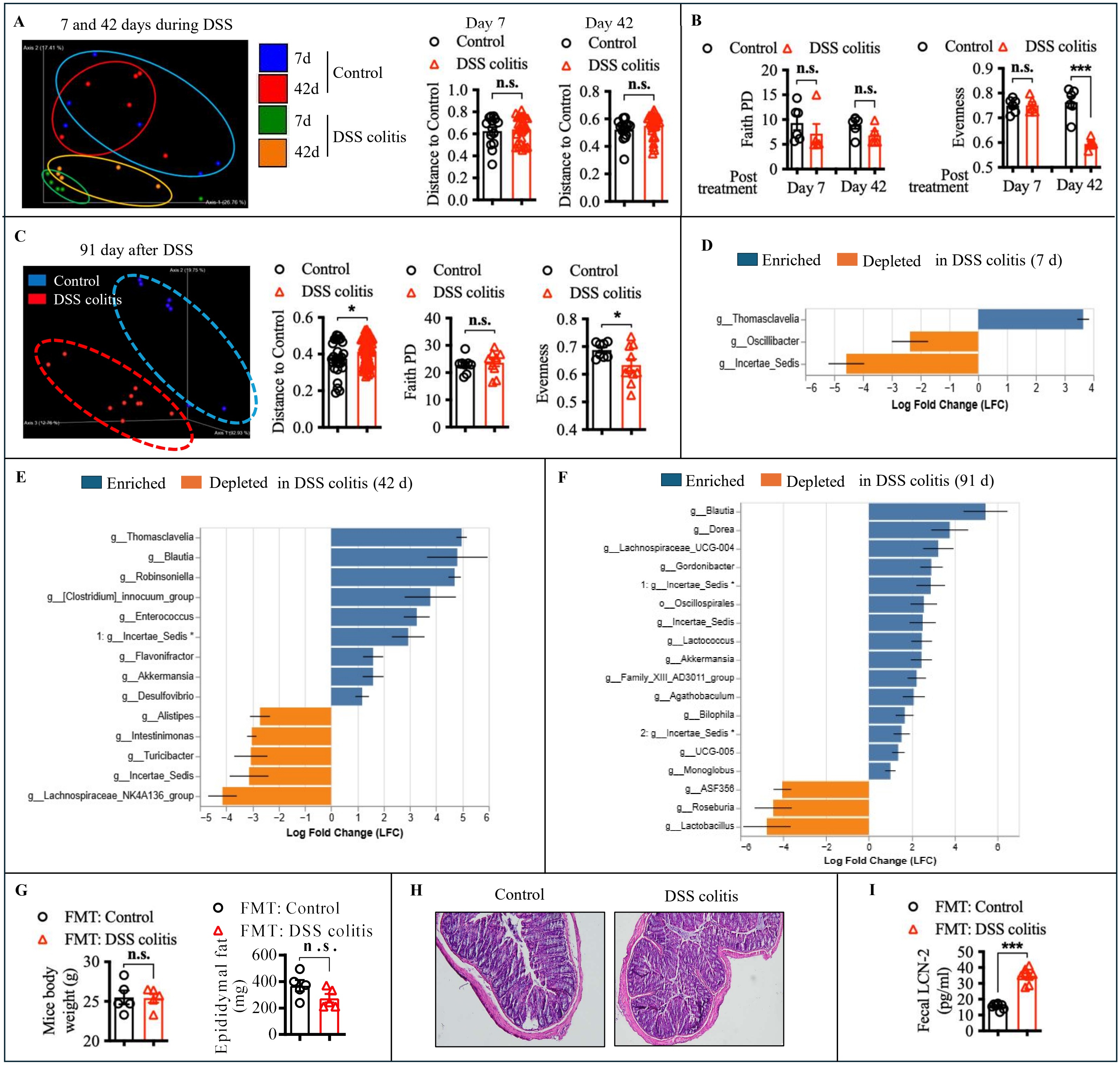


**Figure S7. Dysbiosis persists after colitis and is sufficient to transfer intestinal and systemic phenotypes. A** and **B**, Fecal samples were collected from control and DSS-treated mice on days 7 and 42 and subjected to 16S rRNA gene sequencing. Overall microbial community structure was analyzed using unweighted UniFrac principal coordinate analysis (PCoA) (**A**). Microbial richness and evenness (α-diversity) were assessed using the Faith's phylogenetic diversity (PD) index and evenness metrics (**B**). n=6 mice/group. **C**, Fecal samples were collected from control and DSS-treated mice six weeks after DSS withdrawal. Global microbiota composition was evaluated by unweighted UniFrac PCoA, and α-diversity was assessed using the Faith's PD index and evenness. n=8-10 mice/group. **D–F,** LEfSe (linear discriminant analysis effect size) analysis was performed to identify differentially abundant bacterial genera between DSS-induced colitis mice and control mice at the indicated time points. **G** through I, Germ-free mice received fecal microbiota transplants (FMT) from either healthy control donors or DSS-induced colitis donors. Two months after transplantation, mice were euthanized, and body weight and epididymal fat mass were recorded (**G**). n=5 mice/group. Colonic histopathology was assessed by H&E staining (**H**), and fecal lipocalin-2 (LCN2) levels were measured by ELISA (**I**). Data are presented as mean ± S.E.M. with statistical significance indicated as follows: * $p<0.05$, ** $p<0.01$, *** $p<0.001$.

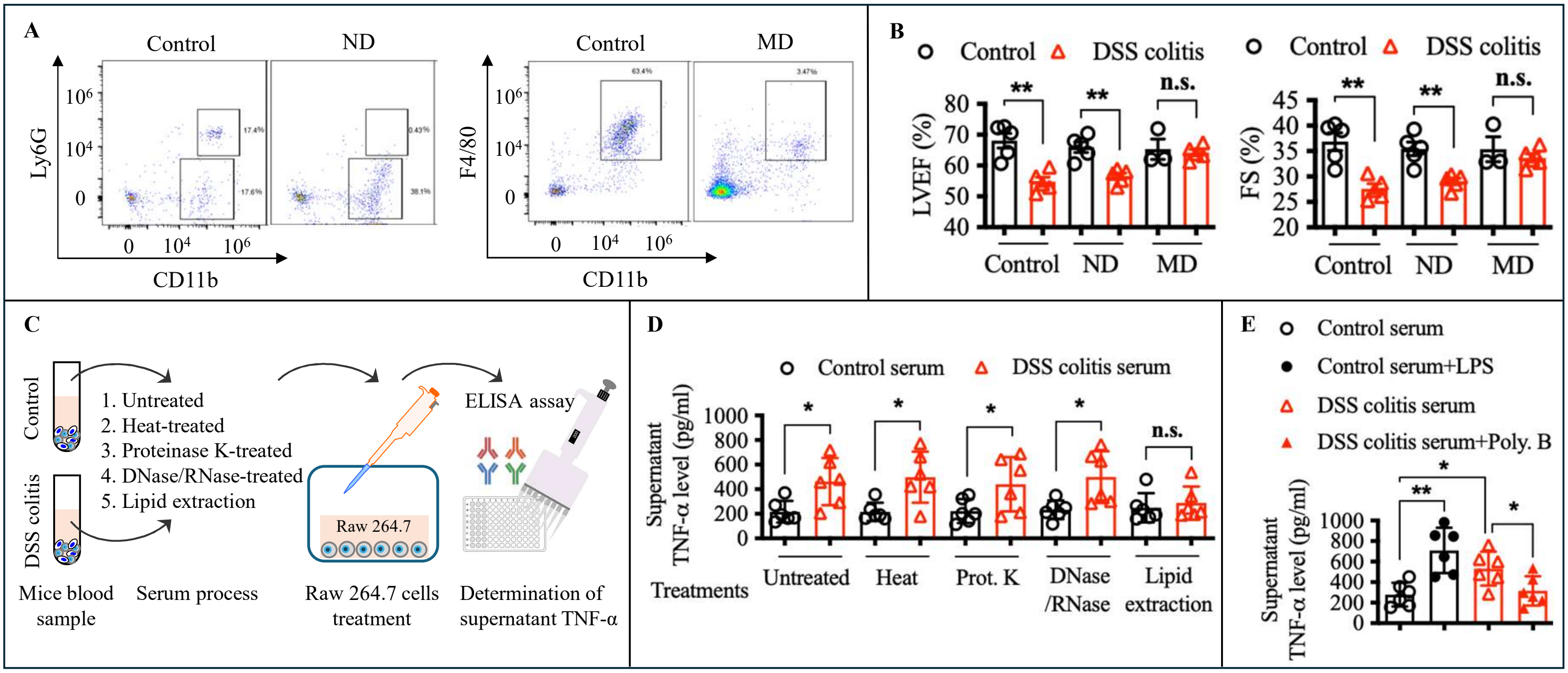


**Figure S8. Identification of LPS as the active circulating factor and requirement of hematopoietic TLR4 signaling.** A. Representative flow cytometry plots showing neutrophils in the heart after anti-Ly6G treatment (ND) or isotype control treatment, and macrophages in the heart after anti-CSF1R plus clodronate liposome treatment (MD) or PBS control treatment. B. Quantification of left ventricular ejection fraction (LVEF) and fractional shortening (FS) in mice after neutrophil or macrophage depletion at 28 days post-DSS-induced colitis**.** **C-E**, Serum from control and DSS-treated mice were subjected to selective biochemical treatments, including heat inactivation, protein digestion (Proteinase K), nucleic acid degradation (DNase/RNase), and lipid extraction. Treated serum were subsequently applied to RAW264.7 macrophages to assess pro-inflammatory activity *via* TNF-α induction. Schematic overview of the workflow used to identify active circulating factors in DSS-induced colitis (**C**). Supernatant TNF-α level after serum treatment (**D**). n=6 mice/group. **E**, RAW264.7 cells were treated with DSS serum in the presence or absence of polymyxin B. Supernatant TNF-α level after serum treatment. n=6 mice/group.

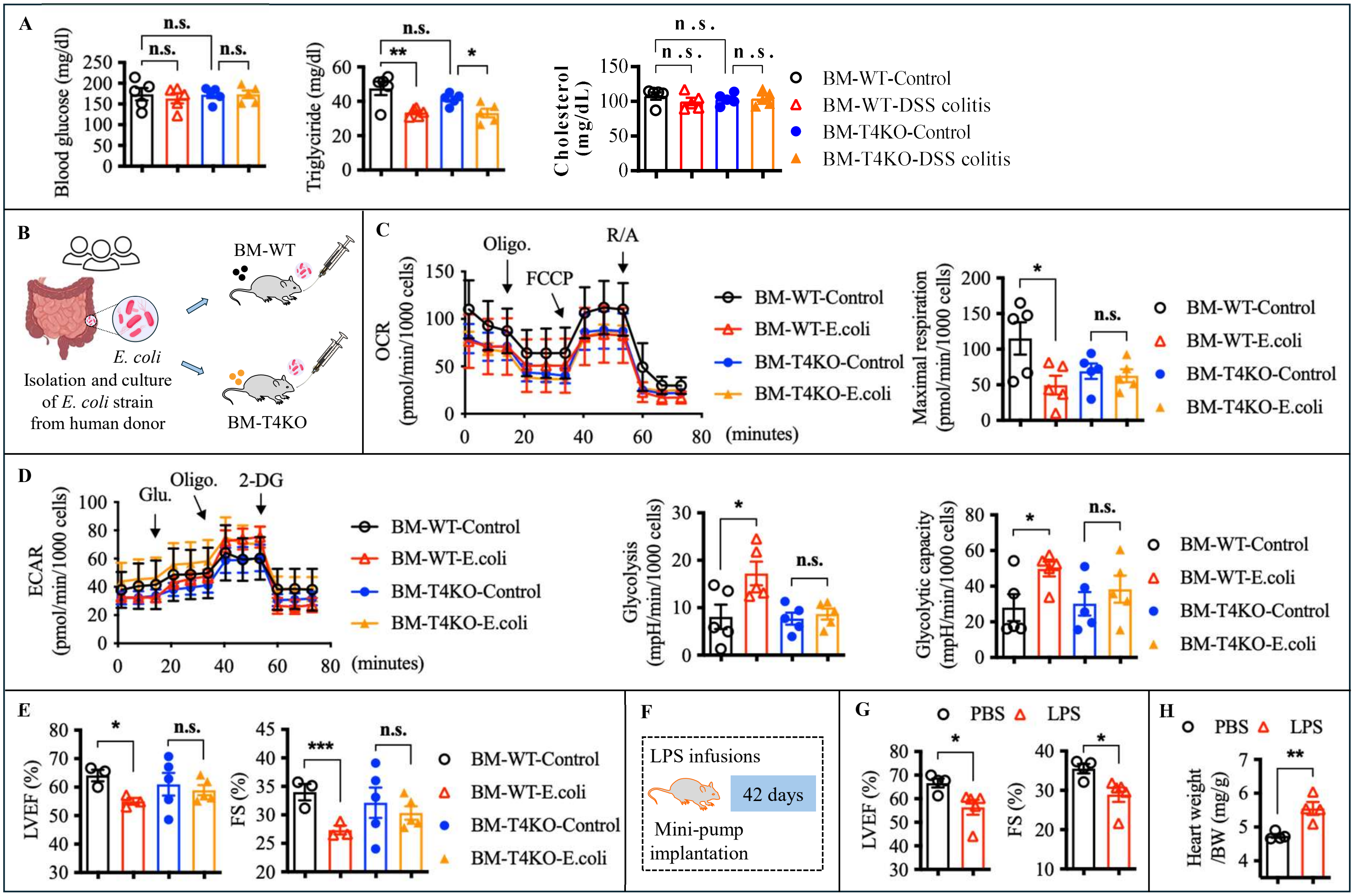


**Figure S9. Hematopoietic TLR4 signaling mediates microbiota- and LPS-induced immune metabolic reprogramming and cardiac dysfunction. A**. Serum levels of glucose, cholesterol, and triglycerides, were assessed in Chimeric BM-WT or BM-T4KO mice (generated as shown in Figure 5B) of control and DSS induced colitis mice at the end of the experiment. **B**, Chimeric BM-WT or BM-T4KO mice were colonized with a commensal *E. coli* strain. **C** through **E**, In chimeric mice, circulating immune cell metabolism was evaluated by Seahorse extracellular flux analysis (**C**and **D**), and cardiac function was assessed by echocardiography (**E**). **F** through **H**, Wild-type mice were chronically infused with LPS using osmotic minipumps (**F**). Cardiac function was assessed by echocardiography (**G**), and cardiac weight was analyzed to evaluate hypertrophy (**H**). n=3-5 mice/group. Data are presented as mean ± S.E.M. with statistical significance indicated as follows: n.s., not significant, * p<0.05, ** p<0.01, *** p<0.001.

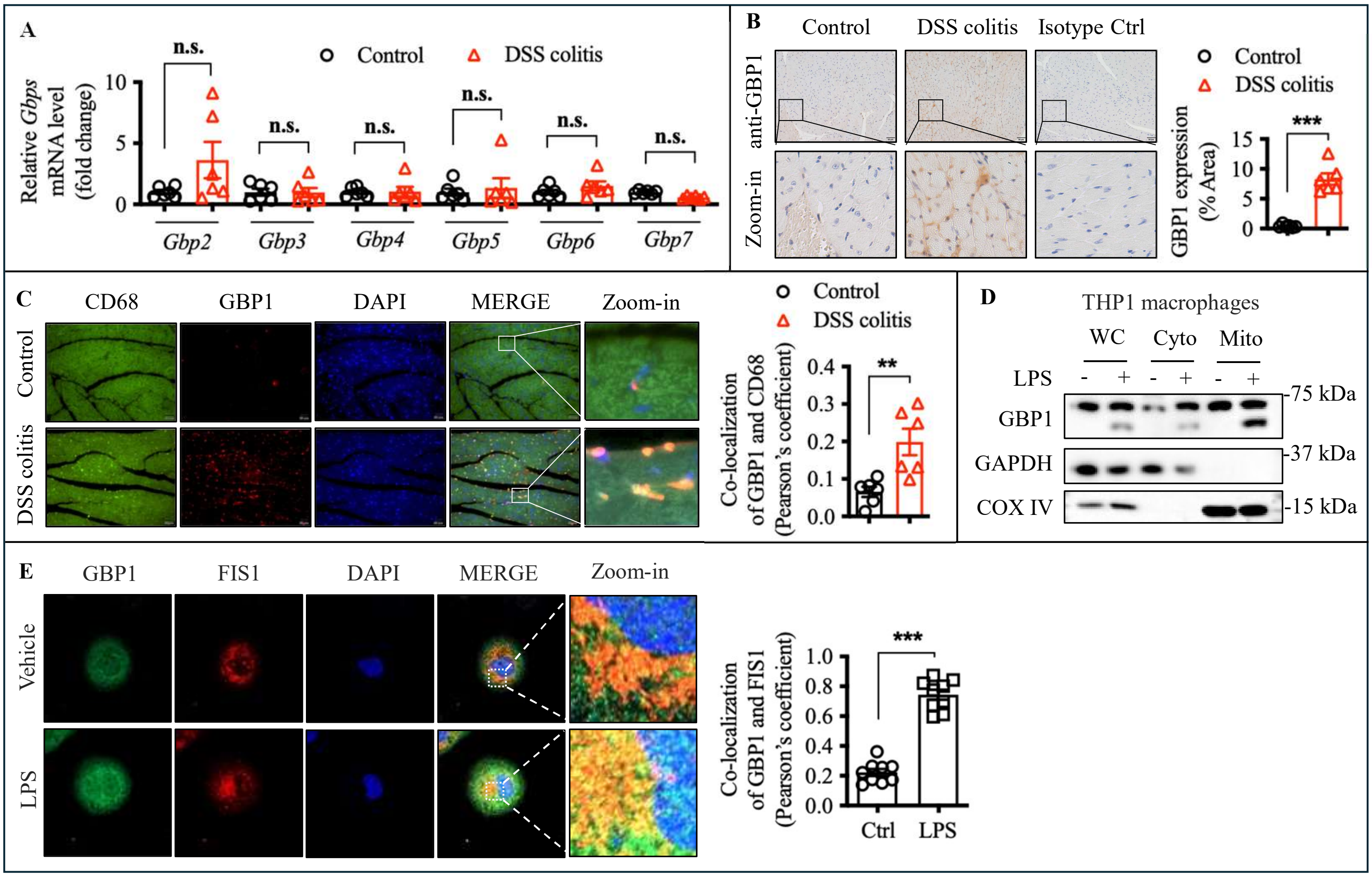


**Figure S10. LPS-TLR4 signaling induces GBP1 expression and mitochondrial localization in immune cells.** **A**, Quantitative RT-PCR analysis of guanylate-binding proteins (GBPs) expression in circulating immune cells at day 42 post-DSS treatment. n=6 mice/group. **B**, GBP1 expression in heart tissue was assessed by immunohistochemistry, and GBP1-positive area was quantified at day 42 post-DSS treatment. n=6 mice/group. **C**, Heart sections were stained for CD68, GBP1, and DAPI, and the colocalization of CD68 with GBP1 was quantified. n=6 mice/group. **D**, Subcellular fractionation of THP-1 monocytes following LPS stimulation, followed by immunoblot analysis of GBP1 in whole-cell, cytosolic, and mitochondrial fractions. **E**, Confocal microscopy of THP-1 cells stained for GBP1 and the mitochondrial marker FIS1 to assess their colocalization after LPS treatment. n=9. Data are presented as mean ± S.E.M. with statistical significance indicated as follows: n.s., not significant, ** $p<0.01$, *** $p<0.001$.

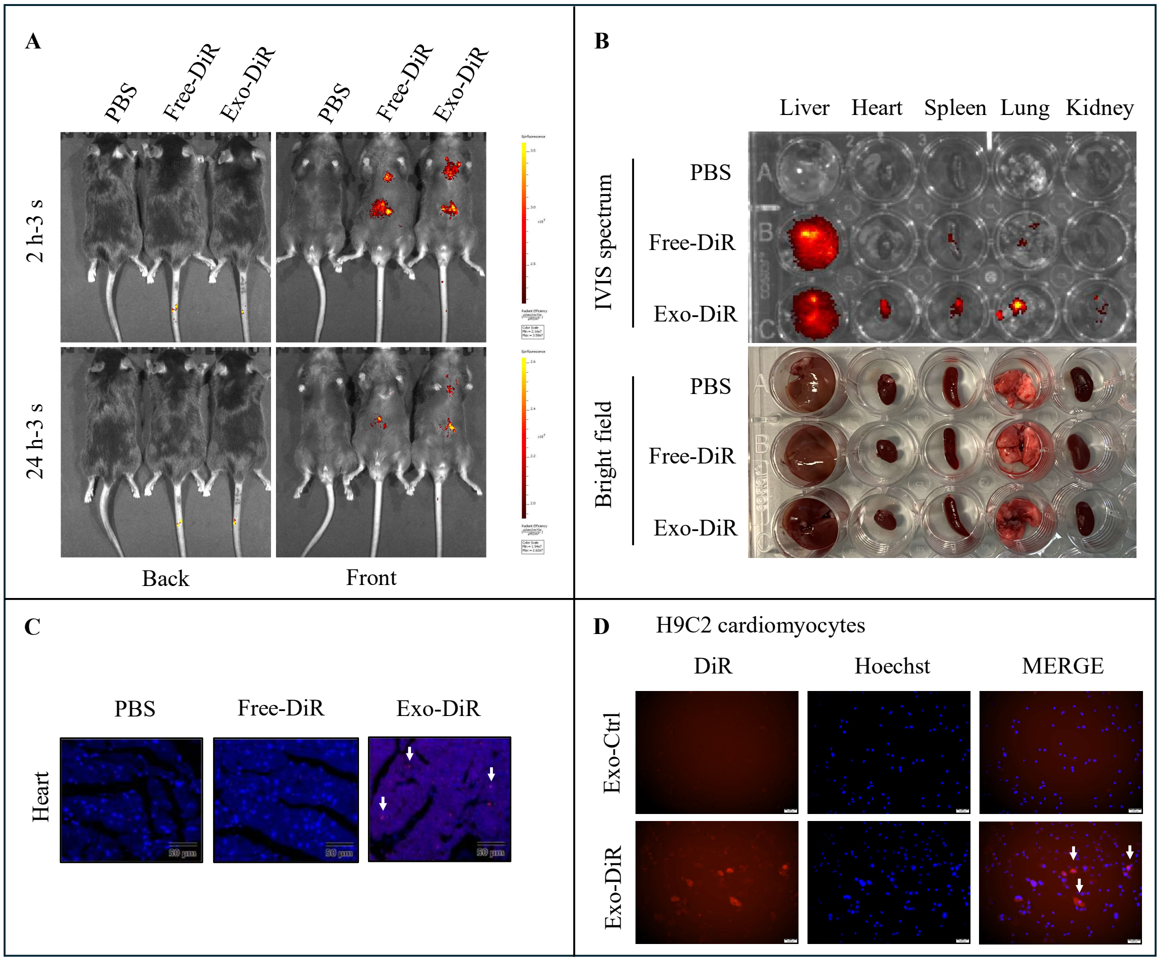


**Figure S11. Systemic biodistribution and cardiomyocyte uptake of circulating exosomes. A** through **C**, DiR-labeled exosomes (Exo-DiR) were administered to mice *via* tail-vein injection. Control mice received an equivalent amount of PBS or free DiR dye (Free-DiR) were processed under identical conditions. *In vivo* fluorescence signals of mice whole body after 2- or 24-h injection (**A**) and *ex vivo* fluorescence images of major organs from mice 24 hours post injection (**B**) were detected by IVIS system, and fluorescence signals (red) in the heart were further confirmed by staining with DAPI (blue) in the frozen sections (**C**). **D**, H9C2 cardiomyocytes were treated with control or DiR-labeled exosomes for 24 h, DiR signals (red) and Hoechst-stained nucleus (blue) were captured by a fluorescent microscopy.

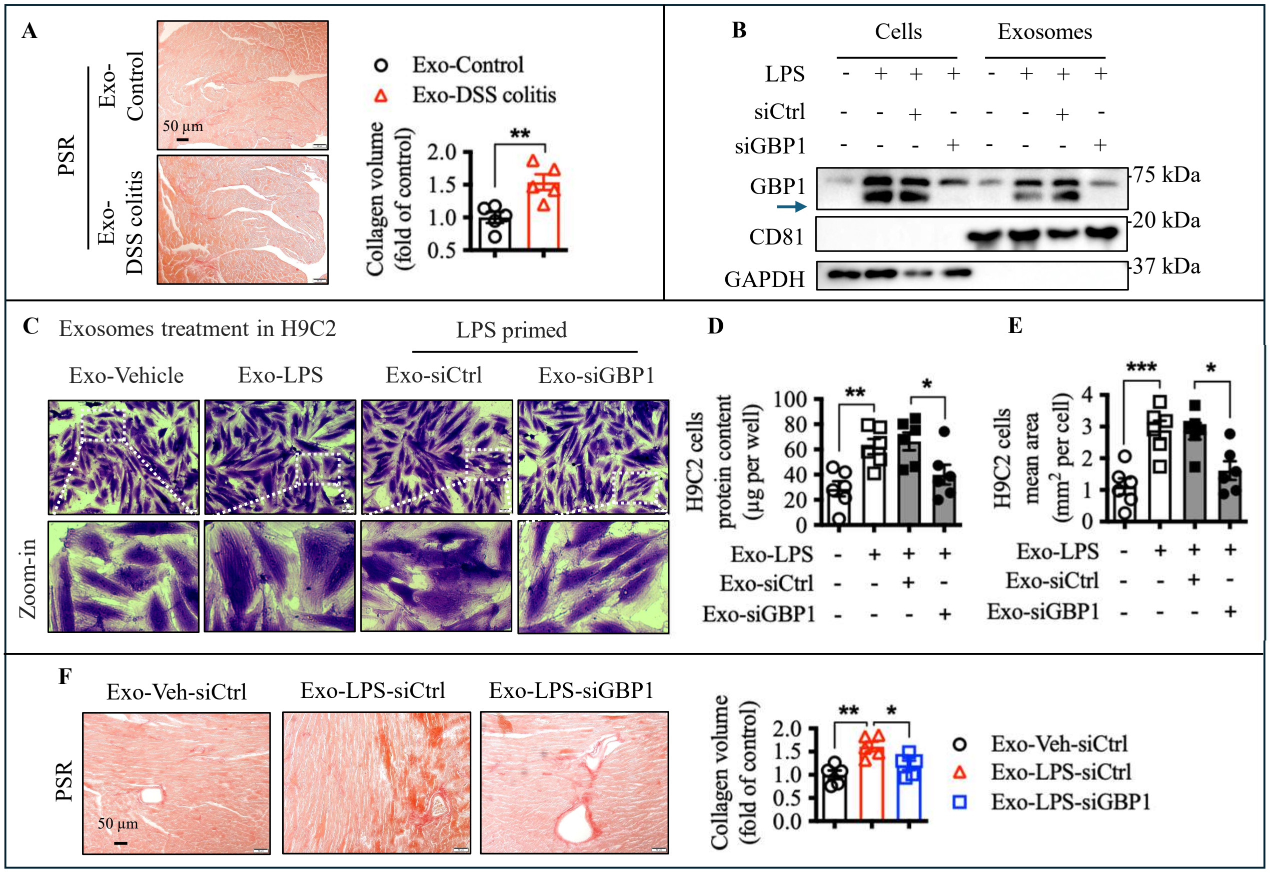


**Figure S12. GBP1-containing exosomes in colitis mediate cardiomyocyte hypertrophy and cardiac fibrosis.** **A**, Wild-type mice were intravenously injected with exosomes isolated from serum of control or DSS-induced colitis mice. Cardiac fibrosis was assessed by Picrosirius Red (PSR) staining, and collagen content was quantified. n=5 mice/group. **B**, THP-1 cells were treated with LPS with or without GBP1 silencing. GBP1 expression in cells and secreted exosomes was analyzed by immunoblotting. **C** through **E**, Exosomes isolated from vehicle- or LPS-treated THP-1 cells, with or without GBP1 silencing, were used to treat H9c2 cardiomyocytes. Cardiomyocyte hypertrophy was assessed by measuring cell surface area, with representative images shown (**C**) and quantitative analyses of total protein content (**D**) and cell size (**E**). n= 6. **F**, Wild-type mice were intravenously injected with exosomes isolated from THP-1 cells treated with vehicle or LPS, with or without GBP1 silencing. Cardiac fibrosis was evaluated by PSR staining, and collagen content was quantified. n=5-6 mice/group. Data are presented as mean ± S.E.M. with statistical significance indicated as follows: * $p<0.05$, ** $p<0.01$, *** $p<0.001$.
